# SVCollector: Optimized sample selection for cost-efficient long-read population sequencing

**DOI:** 10.1101/2020.08.06.240390

**Authors:** T. Rhyker Ranallo-Benavidez, Zachary Lemmon, Sebastian Soyk, Sergey Aganezov, William J. Salerno, Rajiv C. McCoy, Zachary B. Lippman, Michael C. Schatz, Fritz J. Sedlazeck

## Abstract

An increasingly important scenario in population genetics is when a large cohort has been genotyped using a low-resolution approach (e.g. microarrays, exome capture, short-read WGS), from which a few individuals are selected for resequencing using a more comprehensive approach, especially long-read sequencing. The subset of individuals selected should ensure that the captured genetic diversity is fully representative and includes variants across all subpopulations. For example, human variation has historically been focused on individuals with European ancestry, but this represents a small fraction of the overall diversity.

To address this goal, SVCollector (https://github.com/fritzsedlazeck/SVCollector) identifies the optimal subset of individuals for resequencing. SVCollector analyzes a population-level VCF file from a low resolution genotyping study. It then computes a ranked list of samples that maximizes the total number of variants present from a subset of a given size. To solve this optimization problem, SVCollector implements a fast greedy heuristic and an exact algorithm using integer linear programming. We apply SVCollector on simulated data, 2504 human genomes from the 1000 Genomes Project, and 3024 genomes from the 3K Rice Genomes Project and show the rankings it computes are more representative than widely used naive strategies. Notably, we show that when selecting an optimal subset of 100 samples in these two cohorts, SVCollector identifies individuals from every subpopulation while naive methods yield an unbalanced selection. Finally, we show the number of variants present in cohorts of different sizes selected using this approach follows a power-law distribution that is naturally related to the population genetic concept of the allele frequency spectrum, allowing us to estimate the diversity present with increasing numbers of samples.

## Introduction

In recent years it has become increasingly clear that structural variants (SVs) play a key role in evolution, diseases, and many other aspects of biology across all organisms (Lupski 2015; Sudmant et al. 2015; Alonge et al. 2020). It is less well known, however, whether the evolutionary forces shaping SV diversity are analogous or distinct from those influencing single nucleotide variants (SNVs). Genome-wide inferences of human evolutionary relationships (The 1000 Genomes Project Consortium 2015) and key population genetic parameters such as *θ* (Watterson 1975), *π* (Nei and WH Li 1979), and Tajima’s D (Tajima 1989) have largely focused on SNVs but not SVs. Similarly, genome-wide scans of human SNV data have revealed positive and/or balancing selection targeting genomic regions including lactase, the ABO blood groop, and the HLA immune complex (Fu 2014), but the role of SVs in human adaptation remains poorly understood.

Performing population genetic research using structural variants will require better methods that identify SVs in a more cost-effective way. Short-read sequencing is currently the most widely-used approach for identifying SVs, although it suffers from limited accuracy (Chaisson et al. 2015; Sedlazeck et al. 2018). Long reads, such as those from PacBio and Oxford Nanopore, provide greater sensitivity and lower false discovery rates, but their higher costs hinder widespread application in large sequencing studies. Another related question with large cohorts is how to efficiently validate a large number of SVs from the short read-based calls. Traditional methods such as PCR/Sanger sequencing are costly and labor intensive, necessitating careful consideration of variants and samples to validate for further study. Thus, these methods are often limited to hundreds of SVs that can be validated out of an average of 20,000-23,000 SVs present in an healthy individual (Mahmoud et al. 2019).

Here we present SVCollector (https://github.com/fritzsedlazeck/SVCollector), an open-source method (MIT license) to optimally rank and select samples based on variants that are shared within a large population. By default, the optimal ranking strives to capture as much genetic population diversity as possible in a fixed number of samples. As a consequence of this approach, the selected samples will include most common variants plus as many rare and private variants as possible. Alternatively, it can optimize the selection by weighting the variants by their allele frequency, which enriches for common variants in the population. Together, SVCollector allows for both a more cost-efficient way to validate a large number of common SVs, along with an improved re-sequencing approach to discover SVs that were initially missed by short-read sequencing. Naive methods to select samples include picking a random selection or picking the samples which individually have the most variants. These methods do not account for the fact that variants may be shared across multiple samples in the selection. Instead, SVCollector uses a greedy approach to identify a set that collectively spans as many variants as possible.

In the analysis, SVCollector reports the cumulative number of distinct variants present for each individual selected. We show that the number of distinct variants follows a power-law distribution, allowing SVCollector to accurately extrapolate to even larger collections of genomes. After fitting a distinct power-law curve for each of the 26 subpopulations in the 1000 Genomes Project, we estimate the number of individuals that would need to be sequenced in order to obtain a given fraction of the total population-specific diversity. This shows that, minimally, many thousands of human genomes need to be sequenced in order to capture the majority of human variants, especially those with African ancestry.

## Results

We assessed the results of SVCollector based on simulated data (**Supplemental Note 1**, **Supplemental Figures 1–2**) and two large short-read sequencing projects involving 2,504, and 3,024 samples each (see **Figure 1**). For each cohort, we focused on selecting an optimal set of 100 diverse samples. Crucially, using SVCollector, the individuals that are identified span all subpopulations, whereas the commonly used ”topN” approach concentrates the selection in a few subpopulations (see below). For all cohorts, the runtime and memory requirements were minimal. For example, for the 1000 Genomes VCF file of 2,504 samples over 66,555 distinct SVs (Sudmant et al. 2015), SVCollector computed the top 100 samples in 67 seconds using 1.7MB memory. Each of the modes had a similar runtime and RAM requirements.

### Sample selection based on SVs from 2,504 human genomes

We assessed SVCollector based on 2,504 human genomes from the 1000 Genomes Project (Sudmant et al. 2015). For our analyses, we used the phase 3 variant callset (The 1000 Genomes Project Consortium 2015) for chromosomes 1 through 22 with all children removed. **Figure 1** shows a summary of the results and **Supplemental Note 2** and **Supplemental Table 1** list the details. We first investigated the distribution of the 100 samples selected by SVCollector across the 5 superpopulations (**Supplemental Table 2**). Notably, the widely used naive *topN* approach selects 99 African samples and 1 American sample, while SVCollector’s optimal *greedy* approach covers all 5 superpopulations containing 57 African, 14 East Asian, 14 South Asian, 8 American, and 7 European samples and represents 25 of the 26 subpopulations, excluding only GBR (**Figures 1B-C**). The *topN* approach oversamples from the African superpopulation since it has a greater number of SVs than the other superpopulations (**Figure 1A**).

**Figure 1:**
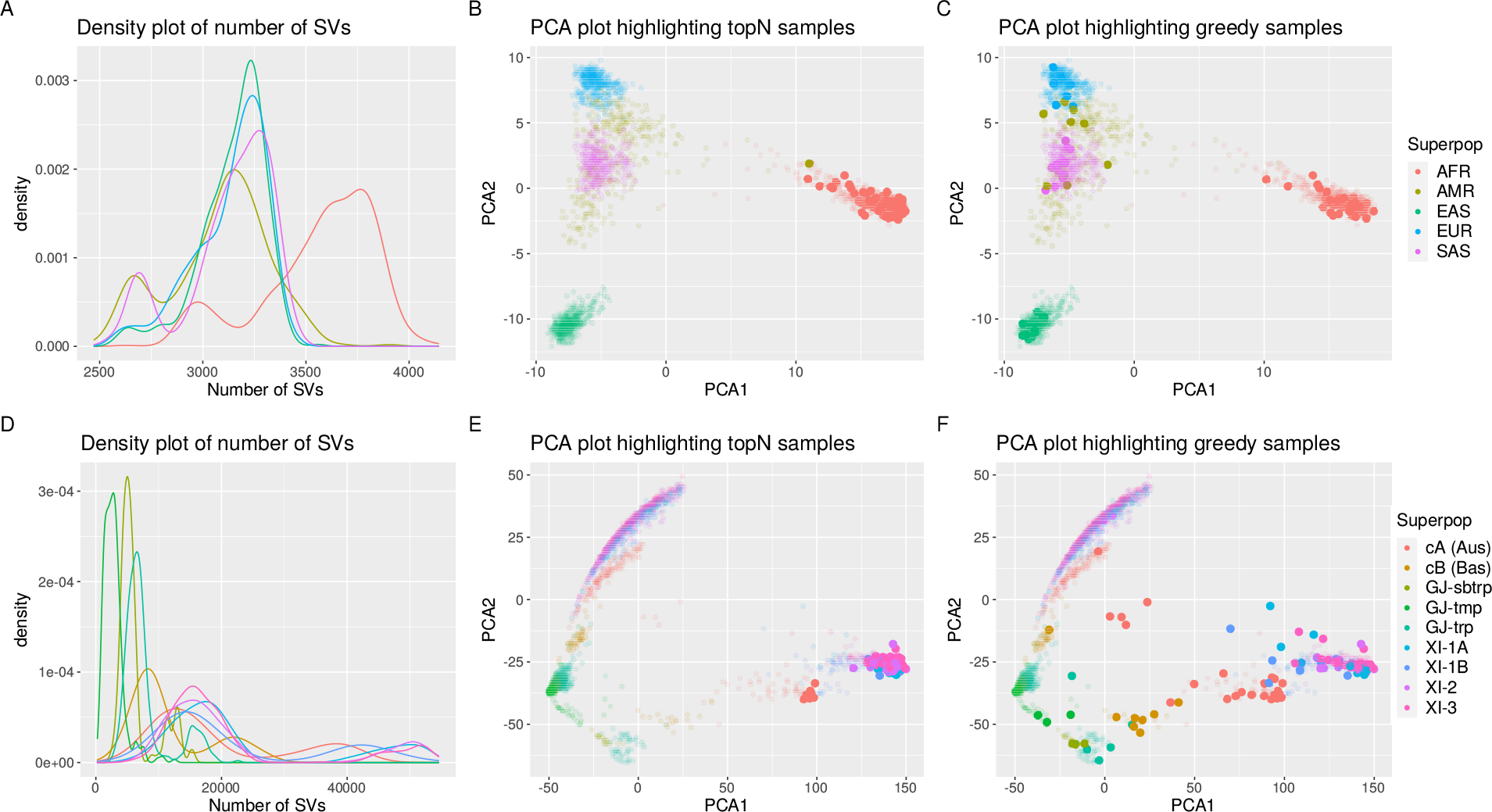
**A)** Density plot of the number of SVs reported per person for each of the five superpopulations in the 1000 Genomes Project. The variants include all non-reference alleles, with both homozygous and heterozygous variants considered equally. The peak for the African (AFR) superpopulation occurs around 3,750 SVs per person, while the peaks for the other superpopulations occur around 3,250 SVs per person. **B-C)** PCA plots of the 1000 Genome Project samples colored by superpopulation highlighting the 100 most diverse samples chosen by the *topN* or *greedy* approaches respectively. The *topN* method oversamples from superpopulations with a greater number of SVs, while the *greedy* method picks a representative sampling across all the superpopulations. **D)** Density plot of the number of SVs per sample for each of the nine populations in the 3K Rice Genomes Project. **E-F)** PCA plots of the 3K Rice Genome Project samples constructed by using variants with an allele frequency greater than 5%. The PCA plots are colored by population and highlight the 100 most diverse samples chosen by the *topN* or *greedy* approaches respectively. The *topN* method oversamples from populations with a greater number of SVs, while the *greedy* method picks a representative sampling across all the populations.

We next investigated the fraction of SVs covered by the 100 samples selected by SVCollector. We compared SVCollector’s *greedy* method to the more widely used *topN* method, to a *random* method, and to the exact algorithm using integer linear programming (*ILP*) (**Figure 2A**). The *ILP* approach allowed us to establish a ground truth so that we could assess the accuracy of the much faster greedy heuristic (**Methods**). The *random* selection was run 100 times per cohort, and we report a boxplot of the percent of SVs identified. The *ILP* solution was run for 24 hours and the best solution at this time was chosen. SVCollector’s fast *greedy* approach (20.43% of SVs) slightly outperforms the widely used naive *topN* approach (19.47% of SVs) and equals the ILP solution (20.43% of SVs) when investigating the top 10 ranked samples in terms of SVs captured. However, when extending the selection to 100 genomes, the *greedy* approach (41.65% of SVs) more substantially outperforms the *topN* approach (35.75% of SVs) and only slightly underperforms the ILP solution (41.75% of SVs) by 74 SVs. Across all the values we tested (k = 5, 6, 10, 12, 15, 16, 20, 30, 40, 50, 60, 70, 80, 90, 100, 120, 140, 160, 180, 200) we find that the greedy approach takes only seconds to run and underperforms the ILP solution by at most 74 variants (0.11% of SVs).

**Figure 2:**
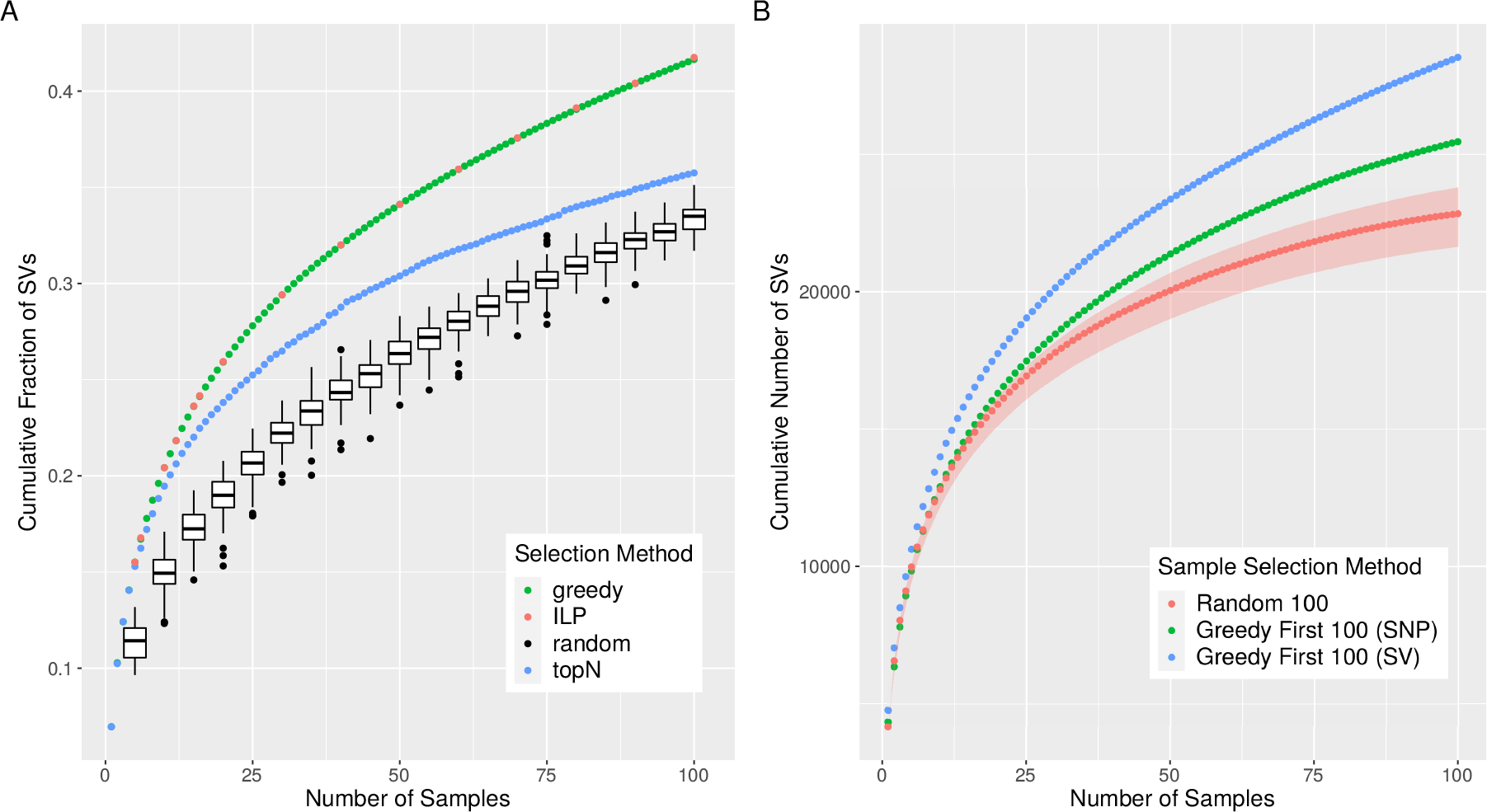
**A)** Cumulative fraction of SVs covered for a given number of samples chosen by the *ILP, greedy, topN*, and *random* approaches. SVCollector’s *greedy* approach approximates the true *ILP* solution and exceeds the *topN* and *random* approaches at recovering unique SVs. **B)** Number of SVs covered using three sample selection methods. In red is the median number of SVs covered over 100 trials of a random sample of 100 individuals. The red ribbon comprises the minimum and maximum number of SVs covered over the 100 trials. In green is the number of SVs covered using the 100 best ranked (greedy) individuals from the SNV data, and in blue is the number of SVs covered using the 100 best ranked individuals (greedy) from the SV data. Data is from the 1000 Genomes Project.

By default, SVCollector maximizes the count of distinct variants, without taking into account the allele frequency of the variants. This often leads to an enrichment of rare or private variants in the identified set, potentially at the expense of capturing more common variants. However, SVCollector can also be run in a mode that takes allele frequency into account. In this mode, SVCollector also uses a greedy approach but optimizes for common variants in the population by weighting variants by their observed allele frequency. We assessed SVCollector in this allele frequency mode to choose the most diverse set of 10 samples in the 1000 Genomes Project. In the allele frequency mode, SVCollector selects samples that cover 92.47% of the total weighted SV diversity, while in the normal mode SVCollector selects samples that cover 91.25%. We also compared the two modes when choosing the most diverse set of 100 samples. In the allele frequency mode, SVCollector selects samples that cover 98.96% of the total weighted SV diversity, while in the normal mode that samples selected cover 98.89%. Furthermore, SVCollector chooses 57 African, 14 East Asian, 13 European, 12 South Asian, and 10 American samples. Thus, even in the allele frequency mode SVCollector chooses a representative selection of samples across all subpopulations.

### Sample selection based on SNVs from 2,504 human genomes

Next, we investigated the relationship between SNVs and SVs, especially to measure if SNV calls can be utilized as an approximation of SV diversity. For this, we used the 1000 Genomes Project data and compared three different methods for picking a sample of 100 individuals to optimize the total number of SVs covered. Overall we find that SVCollector is effective at optimizing sample selection to maximize the number of distinct SVs, even in the absence of SV calls (**Figure 2B**).

First we analyzed 100 trials each consisting of 100 randomly picked individuals. Out of the 100 trials, the SVs covered ranged from 21,627 to 23,792 with a median of 22,839.5 SV. Next, we ran SVCollector in the greedy mode on the SNV data from the 1000 Genomes Project and picked the best ranked 100 individuals. The number of SVs contained in this sample was 25,459. Comparing this to the greedy selection of SVCollector based on SVs resulted in only 3,070 fewer SVs. Thus, selecting the best ranked individuals from the SNV data improves over a random sample, and approaches the upper limit of SVs covered.

### Sample selection based on SVs from 3,024 rice genomes

We also assessed SVCollector based on 3,024 genomes from the 3K Rice Genomes Project (The 3,000 rice genomes project 2014). **Figure 1** summarizes the results and **Supplemental Table 3** lists the details. We first investigated the sampling of the 100 samples selected by SVCollector across the populations, using the 2,223 samples that can be confidently classified into one of the nine populations (W Wang et al. 2018) (**Supplemental Table 4**). Notably, the topN approach selects 42 XI-3, 31 XI-2, 16 XI-1A, 7 cA, and 4 XI-1B samples, but no cB, GJ-subtrp, GJ-tmp, or GJ-trp samples. The greedy approach on the other hand selects a representative sample consisting of all nine subpopulations (24 XI-3, 23 cA, 17 XI-2, 9 XI-1A, 8 cB, 8 XI-1B, 5 GJ-trp, 3 GJ-subtrp, 3 GJ-tmp) (**Figures 1E-F**). The *topN* approach oversamples the XI-3, XI-2, and XI-1A populations since they have a greater number of SVs than the other populations (**Figure 1D**). We next investigated the fraction of SVs covered by the 100 samples selected by SVCollector. SVCollector’s *greedy* approach (19.2%) outperformed the *topN* approach (17.0%) when investigating the first 10 ranked samples. When extending the selection to 100 genomes, the *greedy* approach (45.4%) outperforms the *topN* approach (37.8%).

### New population diversity metrics

Next, we examined the distribution in the number of distinct variants present for each individual selected by SVCollector and found that it follows a power-law distribution (**Methods**). Based on this result, we introduce two new population diversity metrics corresponding to the coefficients *α* and *β* of the best fit power-law curve. Specifically, *α* measures the population mutation rate and *β* measures the extent to which variants are shared across individuals in a population. A larger α value corresponds to a higher mutation rate, while a smaller *α* value corresponds to a lower mutation rate. A less negative *β* (i.e. closer to 0) corresponds to a relative excess of rare variants in the population, while a more negative *β* corresponds to a relative lack of rare variants in the population. Thus, we would expect more genetically heterogeneous populations to have a less negative *β* and more homogeneous populations to have a more negative *β*.

To connect these metrics to well-established theory, we compare our two new population diversity metrics to previously existing metrics. Specifically, Watterson’s *θ* and Tajima’s D are summary statistics derived from the allele frequency spectrum (Fisher 1931; Wright 1938). The allele frequency spectrum considers counts of the number of samples possessing each variant. SVCollector instead considers counts of the number of variants contained within each sample. As we show by using the counts of the number of variants, it is straightforward to extrapolate the number of variants we would expect to see as the number of individuals in the sample is increased.

For each of the 26 subpopulations in the 1000 Genomes data, we calculated a value of *α*, *β*, Watterson’s *θ*, and Tajima’s D over the autosomes. We calculated one set of values on the SNV data and another on the SV data (**Supplemental Table 5**). The program scikit-allel (Miles et al. 2019) was used to determine the values for Watterson’s *θ* and Tajima’s D (**Supplemental Table 6**). From this analysis, we find that the SVCollector population diversity metrics are correlated with these previously existing diversity metrics. We first compared our *α* metric to Watterson’s *θ*, which is used to determine the population mutation rate. We find that *α* is highly correlated with Watterson’s *θ* both using SNV data and using SV data (**Supplemental Figures 4A** and **4C**). Interestingly, the correlations also hold when performing a localized analysis of individual chromosomes, although with varying levels of correlation with *r*^2^ varying from 0.7812 (Chromosome 14) to 0.9026 (Chromosome 19). We next compared our *β* metric to Tajima’s D, which is often used to test for deviations from neutrality or demographic equilibrium. Specifically it compares the mean number of pairwise differences to the number of segregating sites. We find that *β* is highly correlated with Tajima’s D (**Supplemental Figures 4B** and **4D**). We also compared *β* to Tajima’s D calculated on SNVs over each autosome for the 26 subpopulations. We find high correlation in all cases, with *r*^2^ varying from 0.7738 (Chromosome 2) to 0.8683 (Chromosome 21).

To better understand whether the population structure of structural variants within a population is similar to that of small variants, we compared the values of each population metric calculated using SV data for each subpopulation to the values calculated using SNV data for each subpopulation (**Supplemental Figure 5**). We find high correlation between the SV and SNV values for all four population diversity metrics with *r*^2^ values of 0.8941, 0.9387, 0.9932, and 0.9459 for *α*, *β*, Watterson’s *θ*, and Tajima’s D respectively. These results indicate that the population structures of structural variants and small variants are highly analogous.

#### Population substructure

We next show that *β* can be used to compare the genetic diversity of subpopulations. After comparing the values of *β* for each of the 26 subpopulations we find *β* is least negative (indicating a relative excess of rare variants) for the seven African populations and most negative (indicating a relative lack of rare variants) for the five East Asian populations, as expected (**Figure 3A**). This comports with previous analyses of intrapopulation diversity showing the African superpopulation to be the most genetically diverse and the East Asian superpopulation to be the least genetically diverse (The 1000 Genomes Project Consortium 2015) as a result of serial founder effects during ancient human dispersal across the globe (Deshpande et al. 2009).

**Figure 3:**
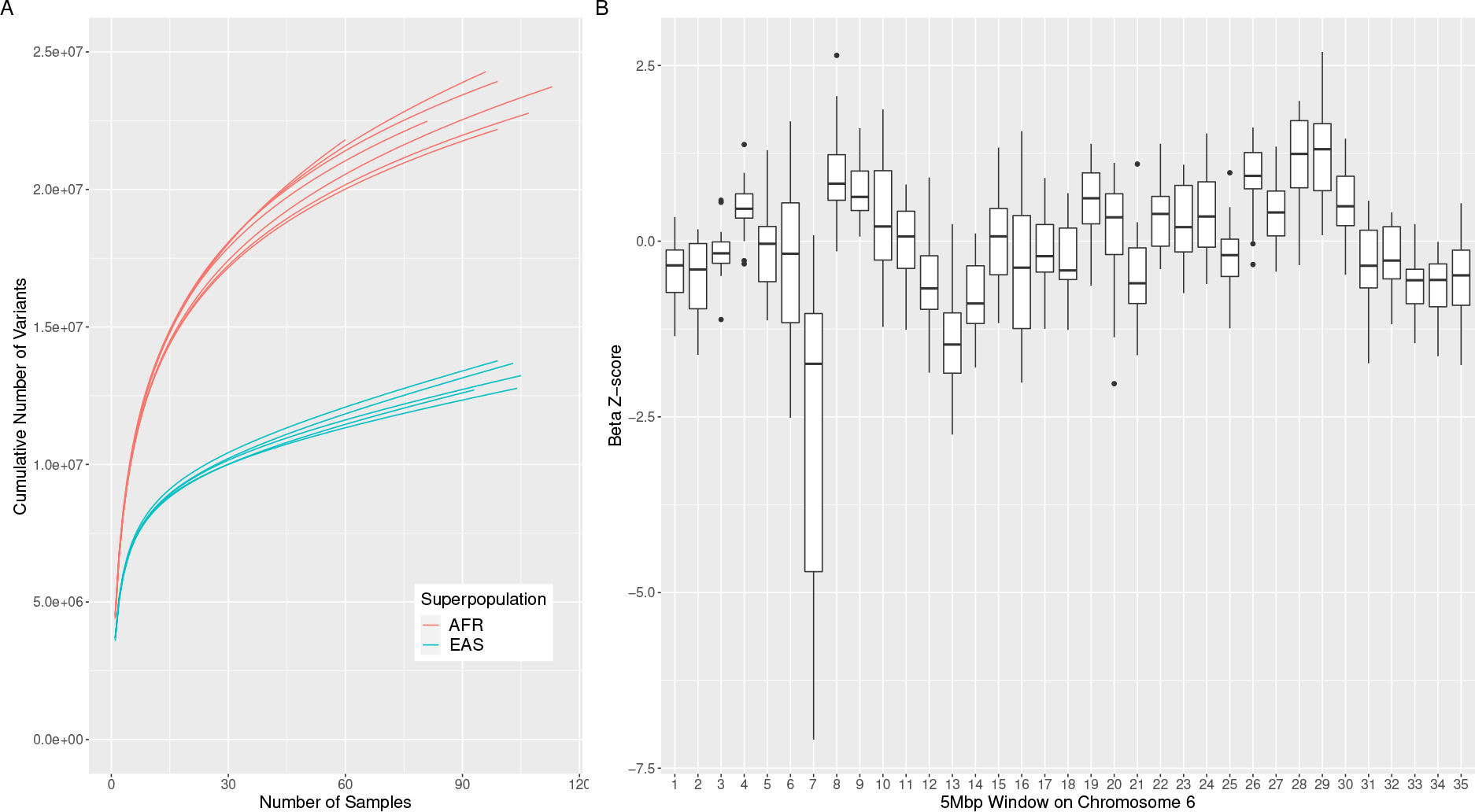
**A)** SVCollector curves for the most and least diverse human subpopulations (SNVs). The seven African subpopulations are the most diverse and the five East Asian subpopulations are the least diverse according to their corresponding *β* values. **B)** Selective sweep on Chromosome 6. For each 5 Mbp window on Chromosome 6, the *β* z-scores for the 26 populations are plotted as a box plot. Window 7 corresponds to the HLA region and shows a strong signal for selective pressure acting in this region.

#### Signatures of selection

We next show that *β* can be used to find regions of the genome exhibiting signatures of positive or balancing selection. To perform a genome-wide scan for such signatures, we calculated *β* over small genomic regions (5 Mbp non-overlapping windows). Specifically, for each of the 26 subpopulations we calculated *β* over each window using only the corresponding SNVs. On average, there were 28,760 SNVs per window. Then, for each subpopulation we computed the *β* z-score across all windows to allow for comparisons across the different subpopulations. For each window, we then constructed a boxplot of the *β* z-scores across the 26 subpopulatons. We find that for the 26 subpopulations, the genomic region spanning the HLA immune complex (window 7 on Chromosome 6) has a more negative *β* value than all other regions. This indicates a relative lack of rare variants in this region which is a signal of balancing/diversifying selection (Hughes and Yeager 1998) (**Figure 3B**).

Additionally, *β* can be used to discover regions of the genome targeted by historical local adaptation, whereby positive selection generated strong frequency differences across human populations. For example, we would expect that Northern European populations but not East Asian populations, would exhibit signatures of positive selection targeting the lactase gene (LCT) (Bayless et al. 2017; Bersaglieri et al. 2004). Indeed we find that the British in England and Scotland (GBR) population has the most negative *β* z-score (−0.920) for this region, while none of the East Asian populations have a negative *β* z-score. The limited number of SVs in the 1000 Genomes dataset, with an average of only 26 SVs per 5 Mbp window, limits similar selective sweep analyses using SVs.

### Extrapolations

Finally, we perform extrapolations of the best fit power-law curves to determine the extent to which the human pan-genome is open and provide lower bounds on how many individuals would need to be sequenced to obtain a given proportion (90%) of the total shared genetic diversity. By shared genetic diversity, we mean all the variants that are either partially or entirely shared by the individuals in the population and excluding singleton variants found only in a single individual. If every individual brings a positive number of singleton variants, it would be impossible to entirely sequence the pan-genome without sequencing every individual. The shared diversity corresponds to the *α*(*i*)^*β*^ term of the power-law model, which is the term we use to make the extrapolations.

First, we perform subsampling on the entire dataset, fit a best fit curve to each subsample, and extrapolate to the full dataset. We run 100 trials each for subsamples of 10, 25, and 100 random individuals (**Figure 4A-B**). As is expected, the extrapolation from subsampling produces an underestimation in the amount of diversity because the subsample would have to include exactly the most diverse individuals in order for the extrapolated diversity to match the actual diversity on the entire dataset. Consequently, increasing the sample size improves the accuracy of the extrapolation.

**Figure 4:**
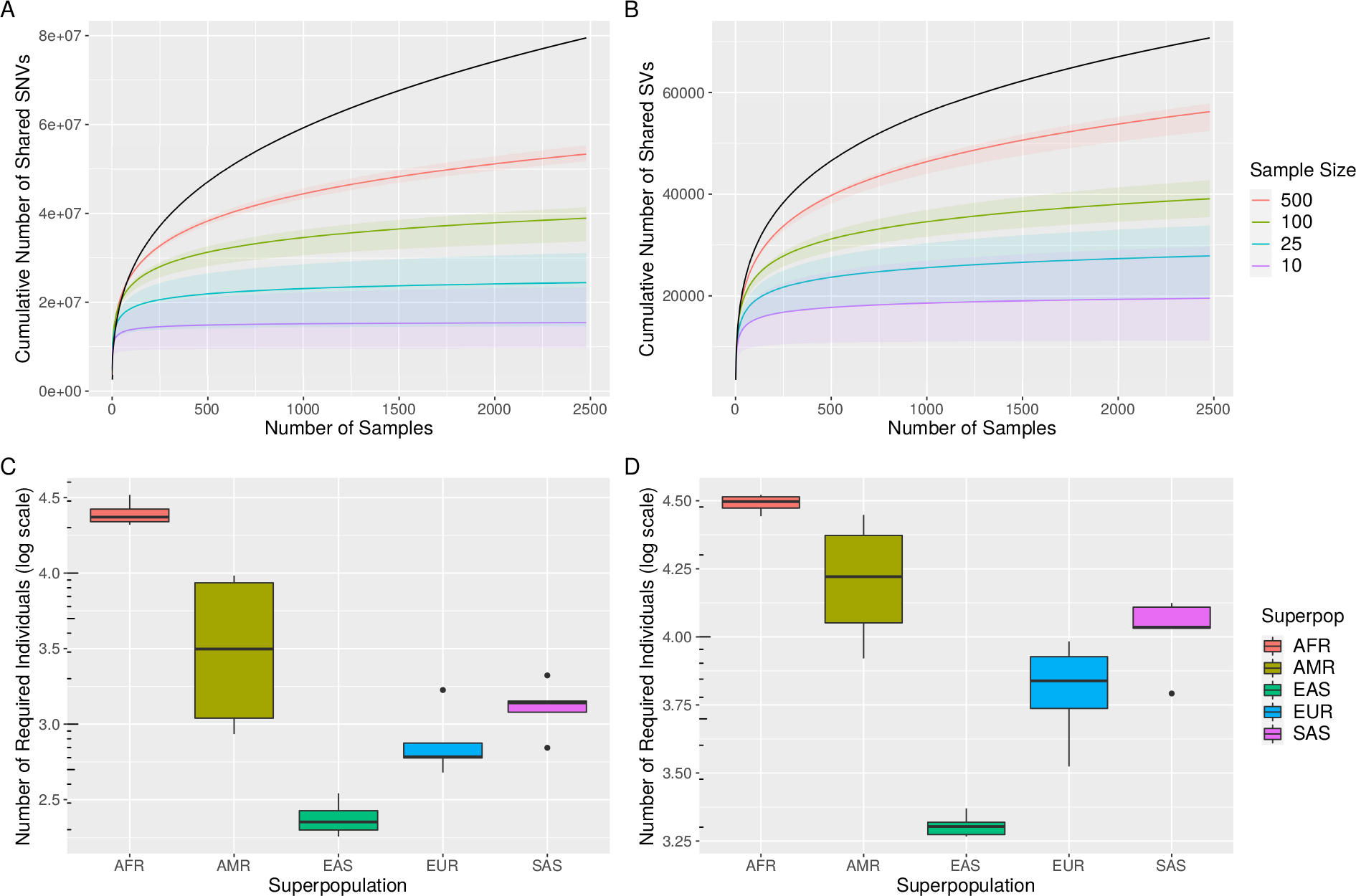
**A-B)** Extrapolating the number of shared variants covered with subsamples. In black is the shared diversity of the full dataset. The lines represent the median number of covered variants over 100 trials of the given sample size. The ribbons represent the minimum and maximum number of covered variants over 100 trials of the given sample size. The left panel is calculated on SNVs while the right panel is calculated on SVs. **C-D)** Boxplots of subpopulation diversity by superpopulation. For each subpopulation, the predicted total amount of shared variants for 100,000 individuals is calculated. Then, the number of sequenced individuals necessary to obtain 90% of this diversity is calculated. Finally, boxplots of the number of required individuals (log scale) are plotted for the corresponding subpopulations in each superpopulation. The left panel is calculated on SNVs while the right panel is calculated on SVs.

Next, for each subpopulation, we extrapolate the best fit power-law curve out to 100,000 individuals to estimate a lower bound on the total number of shared variants. Then, we calculate the number of sequenced individuals necessary to obtain 90% of this diversity. We find that relatively fewer East Asian individuals would need to be sequenced and relatively more African individuals would need to be sequenced (**Figure 4C-D**). For example, only 180 Chinese Dai (CDX) individuals are needed to capture 90% of the shared SNVs of 100,000 CDX individuals, but at least 32,978 Luhya (LWK) individuals are needed to capture 90% of the shared SNVs of 100,000 LWK individuals. Since these estimates are lower bounds, many more individuals need to be sequenced in order to fully capture the diversity of human variants but expect the relative amount of diversity between the subpopulations to remain.

## Discussion

SVCollector is a fast and powerful method to quantitatively and optimally select samples for long-read resequencing or optical mapping based on their genomic variation (SNV and or SV) shared in the population. SVCollector’s *greedy* mode substantially outperforms the commonly used *topN* or *random* selections both in the representativeness across populations and in the number of SVs captured. These gains will in turn translate to cost savings for resequencing and validation experiments. Indeed, SVCollector has already been applied to larger sequencing projects such as a detailed study of 11 human genomes and 100 tomato genomes sequenced with long reads (Shafin et al. 2020; Alonge et al. 2020).

Importantly, we found that SNV variant callsets can be used to choose a sample of individuals that maximizes the number of distinct SVs. In the 1000 Genomes dataset, this method results in only 3,070 fewer SVs captured than when using the SV variant callset directly. Additionally, in the human datasets, female samples often contributed more SVs than male samples because of the extra heterozygous SVs on the X chromosome. Depending on the application, researchers may want to exclude the sex chromosomes prior to analysis as we did in our analyses.

The two new population diversity metrics we introduced, *α* and *β*, can be used to quantify the frequency distribution of SV as well as scan for genomic regions exhibiting signatures of historical selection. *α* and *β* can be easily calculated from multi-sample VCF files, which will allow researchers to gain important insights about understudied populations across species. Unfortunately, the relatively small number of SVs in the 1000 Genomes callset limits similar analyses for human SVs. Furthermore, the values of *α*, *β*, Watterson’s *θ*, and Tajima’s *D* calculated on SV data are highly correlated with the values calculated on SNV data. This helps explain why SVCollector is effective at optimizing sample selection even in the absence of SV calls. Finally, we show that the human pan-genome is very diverse, and that capturing 90% of the total shared diversity would require the sequencing of many more individuals than has been done in any long-read cohort to date.

It remains challenging to create a population-wide variant callset as the selection clearly depends on the quality of the initial SV callset. Nevertheless, given a low quality (i.e. over representation of false positive SVs) variation callset, SVCollector will likely also over-represent false positives, which will help with the detection and negative validation of these SV calls. In the case when an SV callset is limited in the detection of SVs, SVCollector will still rank the samples, but it is unclear what minimal sensitivity rate would be needed to accurately represent the population.

Overall we showcase a cost-efficient yet comprehensive way to utilize long-read sequencing at population scale. This is particularly important for population projects (CCDG, TOPMed, 1000 Genomes Project, etc.) where genotyping data is available. We show that SVCollector identifies the optimal subset of samples for further examination and at the same time provides population specific insights. Given the current many-fold price differences between long-read sequencing and Illumina sequencing together with the abundance of population studies (exons, arrays, etc.), SVCollector will remain useful for quite some time.

## Methods

SVCollector is implemented in C++, and computes a ranked list and diagnostic plots of the samples listed in a multi-sample VCF file. It uses an iterative approach to minimize the memory footprint, and requires less than 2 MB of RAM even when ranking thousands of samples with tens of thousands of variants each. In the first iteration, it parses the VCF file, counts the total number of variants, and generates a temporary file storing the sample IDs associated with each SV. For subsequent iterations, it reads the temporary file and deletes SVs that were present in the previously selected sample.

SVCollector has two major ranking modes: *topN*, and *greedy*. For the *topN* mode, it picks samples with the largest number of SVs irrespective if the SVs are shared with other samples. For the *greedy* mode, it finds an optimal subset of samples that collectively contain the largest number of distinct variants (see **Figure 5** for an example of the difference between these two modes). Solving this exactly is computationally demanding as it is a version of the well-known NP-hard maximum coverage problem. We implement the following integer linear programming (*ILP*) formulations of the problem to solve it exactly, although this requires an excessively long run time even for relatively small datasets.

**Figure 5:**
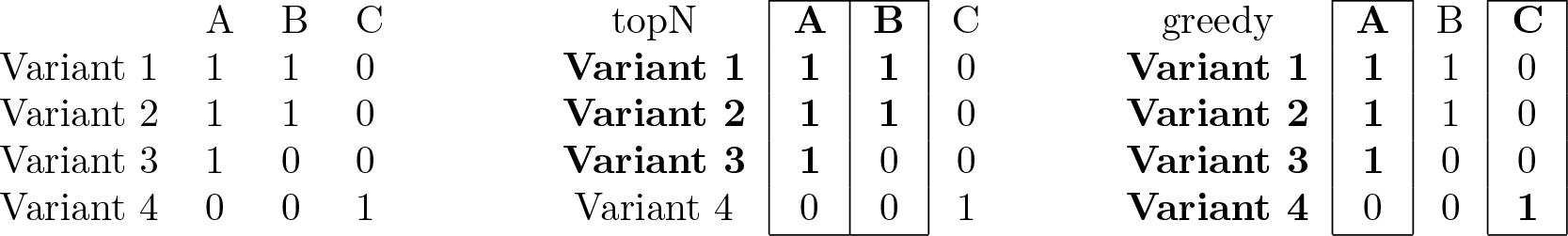
**A)** Presence/absence matrix for three samples and four variants. **B)** When picking the two most diverse samples, the *topN* algorithm selects A and B since they are the individuals with the most number of variants. However, this selection only includes three of the four variants. **C)** The *greedy* algorithm on the other hand selects A and C since it accounts for the fact that the variants covered by B have already been included by A. The *greedy* selection includes all four variants.

The input for the *ILP* maximum coverage optimization problem is represented by an *n* × *m* binary matrix *A* = [*a_i,j_*] ∈ {0,1}^*n*×*m*^, in which every entry *a_i,j_* ∈ {0,1} determines if in sample *i* the variant *j* is present or absent. Given a matrix *A* and 1 ≤ *k* ≤ *n*, we define an optimization problem as a search for a subset *I* ⊆ {1, 2, …, *n*} of samples with |*I*| ≤ *k*, such that the total number of variants present across all samples in *I* is maximized.

We formulate the following *ILP* to solve the aforementioned optimization problem. First, we define the decision variables in our *ILP* formulation:

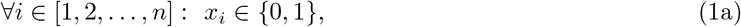

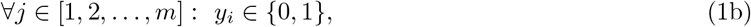

where an *x_i_* encodes whether or not a sample *i* is selected to be present in the problem solution *I*, and a variable *y_j_* encodes whether or not a variable *j* is going to be represented in the problem solution (i.e., present in at least one sample from *I*).

We now define the constraints for the *ILP* formulation. We start with the constraint that ensures that no more than *k* samples are selected:

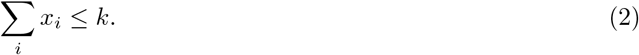

We then define constraints that ensure that when a variable *y_j_* = 1, at least one sample *x_i_* in which variant *j* is present is selected:

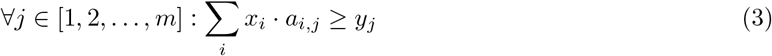

Finally, we define an objective function for our optimization ILP, forcing the maximum number of variants to be represented in the desired solution:

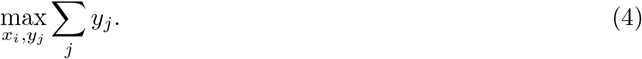

Though these *ILP* formulations solve the maximum coverage problem exactly, they are inefficient. Conversely, the greedy algorithm provides an efficient polynomial time solution that closely approximates the optimal solution (Feige 1998). Consequently, SVCollector uses a greedy approximation that starts with the sample with the largest number of variants, and then iteratively picks the sample containing the largest number variants not yet included in the subset. It also has a *random* mode that mimics an arbitrary selection process, and is helpful for evaluating the diversity of the *topN* or *greedy* approaches. For each mode, SVCollector reports the rank, sample name, its unique contribution of SVs, the cumulative sum of SVs up to the chosen sample, and the cumulative percentage compared to the total number of SVs in the input VCF file.

### Power-law curves

SVCollector creates a diagnostic plot of population diversity where the y-axis is the cumulative count of variants up to the chosen sample and the x-axis is the number of samples. These SVCollector curves (when produced using the *greedy* mode) allow us to visualize the rate at which the cumulative number of variants increases as individuals are optimally added. Indeed, this rate is a function of the genetic diversity of the population under consideration. To see this, consider a population consisting of individuals with a constant positive number of personal variants, but with zero shared variants between each other. In this case, the rate of change in the cumulative number of variants will remain constant as individuals are added. Now consider a population consisting of individuals with shared variants. A higher prevalence of shared variants across individuals will result in a faster decrease in the rate of change of cumulative variants as individuals are added.

Thus, it would be beneficial to model the shape of these curves so that information about the underlying population diversity can be extracted. We have found that these curves are modeled exceptionally well by a power-law distribution (**Figure 6** and **Supplemental Figures 6**-31). The power-law distribution has been found to underlie many natural phenomena, and arises from situations involving a preferential attachment process (i.e. new items are preferentially distributed among individuals according to how many items they already have) (Mitzenmacher 2004).

**Figure 6:**
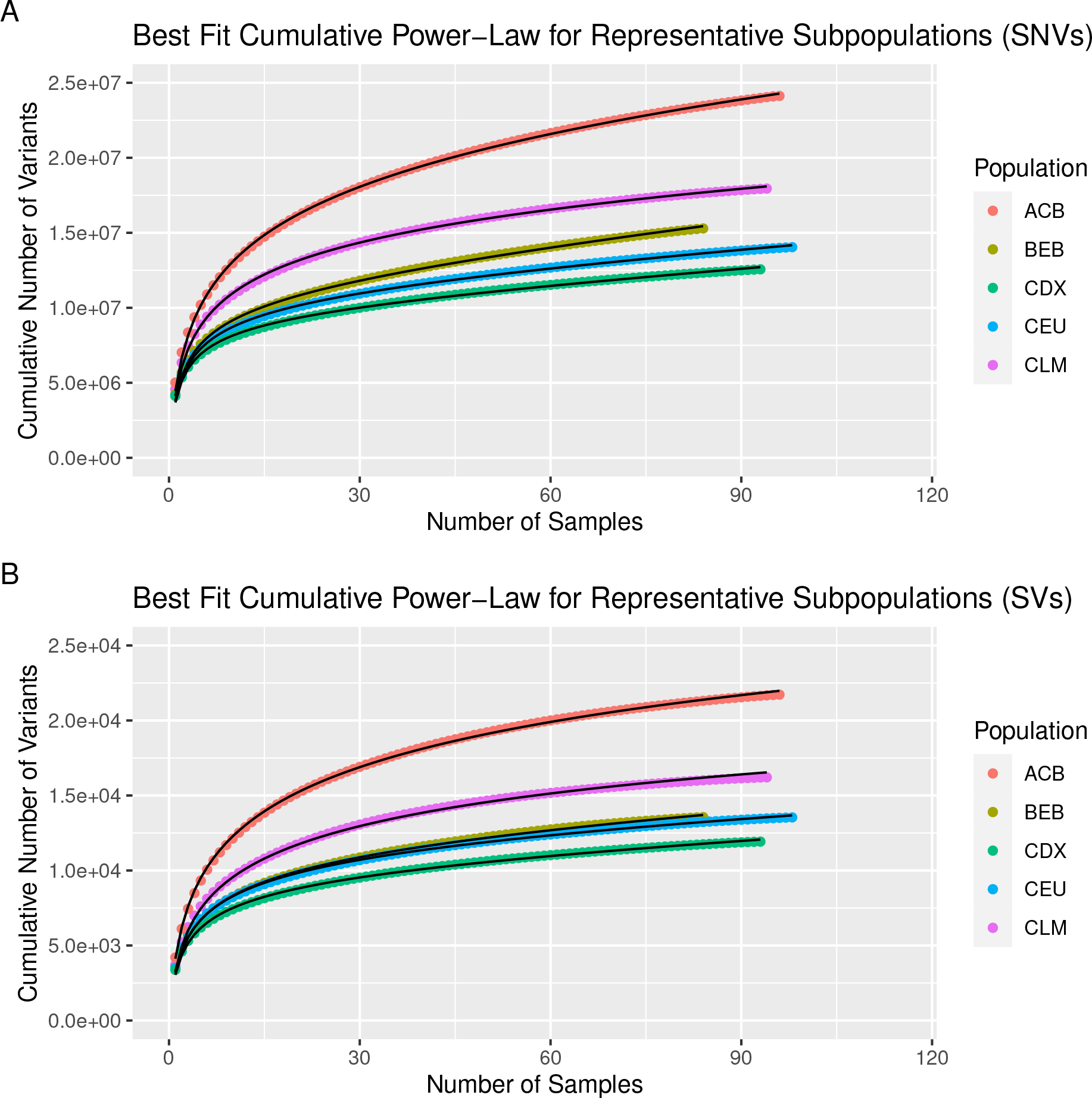
SVCollector curves and best fit cumulative power-law models for **(A)** SNVs and **(B)** SVs. Data is shown for one representative subpopulation for each superpopulation in the 1000 Genomes Project.

Similar analyses of the cumulative population diversity in samples have been performed in the context of determining the extent to which the pan-genome of a bacterial species is open or closed (Medini et al. 2005). In this analysis, the pan-genome includes the “core” genome which is shared among all individuals in a population and a “dispensable” genome which is either partially shared between individuals (accessory genome) or is unique to a single individual (unique genome). Here we define the “shared” genome as the union of the core and accessory genomes. In one study, the pan-genome of *Streptococcus agalactiae* was concluded to be open due to mathematical extrapolation of the plot of the cumulative number of genes present versus number of strains added (Tettelin et al. 2005).

To model the SVCollector curves, we fit the following equation:

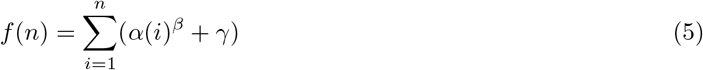

where *n* is the number of individuals included in the subset, *f*(*n*) is the cumulative number of variants present after the *n*-th individual is added, *α* is a population diversity metric that scales with the total number of variants in the population, *β* is a population diversity metric that describes the diversity of the population, and *γ* is a variable that relates to the number of personal variants for each individual. To determine the parameter values of the model that best fit the data, SVCollector uses non-linear optimization. Specifically, the nlsLM function in R is used, which implements the Levenberg-Marquardt algorithm (More 1978) whereby an iterative procedure is performed to update the initial estimate.

## Data Access

The 22 autosome SNV VCF files for the 1KGP project can be downloaded at ftp://ftp.1000genomes.ebi.ac.uk/vol1/ftp/release/20130502/. The 22 autosome SV VCF files for the 1KGP project can be downloaded at ftp://ftp.1000genomes.ebi.ac.uk/vol1/ftp/phase3/integrated_sv_map/. The data used to generate the SV VCF files for the 3K Rice Genome Project Large Structural Variants Dataset can be downloaded at https://snp-seek.irri.org/_download.zul.

## Acknowledgements

This work was supported, in part, by National Science Foundation grants DBI-1350041 and IOS-1732253, and National Institutes of Health grant UM1-HG008898. Part of this research project was conducted using computational resources at the Maryland Advanced Research Computing Center (MARCC).

## Author Contributions

T.R.R.-B. and F.J.S. implemented the SVCollector algorithm. T.R.R.-B. performed mathematical modeling, population diversity metric comparisons, scans for signatures of selection, and extrapolations. S.A. performed the ILP analyses. M.C.S. and F.J.S. supervised the project. All authors contributed to the manuscript.

## Supplemental Note 1. Simulated Population Analysis

For evaluation we wrote a simulation script that generates a multi-sample VCF file with an arbitrary population structure. Briefly, the simulator simulates *F* founder genotypes, that each contain on average *Normal*(*N, M*) variants placed at random along the genome (the initial genome size is fixed at 100,000,000 bp). Then for each founder population, a collection of *Normal*(*S,T*) individual samples are generated at random that contain the original founder variants plus an additional *Normal*(*X, Y*) variants. Consequently, the expected total number of variants in the collection is *F* * *N* + *F* * *S* * *X* variants. If *N* > *S*, then most of the variants will be shared within the population group, and if *S* > *N*, most variants will be unique to that sample. We emphasize this is not designed to simulate realistic pedigrees, but to examine the extremes of high or low levels of sharing among the individuals.

The code can be found at our GitHub page: https://github.com/fritzsedlazeck/SVCollector/tree/master/simul.

The first simulated population (simple10.vcf) has 10 founder genomes with exactly 1000 variants located at random. From each founder genome, 10 samples are simulated that contain the 1000 founder variants plus an additional 100 variants. The SVCollector curve for this sample is presented in **Supplemental Figure 1**. The sharp inflection point for the greedy curve at N=10 illustrates how the code realizes there are 10 founder populations. After these 10 populations have been sampled, the rate at which additional SVs are identified reduces to a much slower rate as these variants are only contained in individual samples.

The simulator was executed like this: `./popsim.pl 10 10 0 1000 0 100 0 > simple10.vcf`

The second simulated population (complex10.vcf) also has 10 founder genomes that each randomly contain *Normal*(500, 250) variants. From each founder population, *Normal*(10, 5) samples are simulated that contain the founder variants, plus an additional *Normal*(500, 250) variants unique to this sample. The SVCollector curve is presented in **Supplemental Figure 2**. Notice that despite having the same number of founder genomes, the curves are substantially different, and lack the inflection point at *N* = 10. This highlights how sample specific variants contribute at a similar level to the founder genotypes.

The simulator was executed like this: `./popsim.pl 10 10 .5 500 .5 500 .5 > complex10.vcf`

**Supplemental Figure 1:**
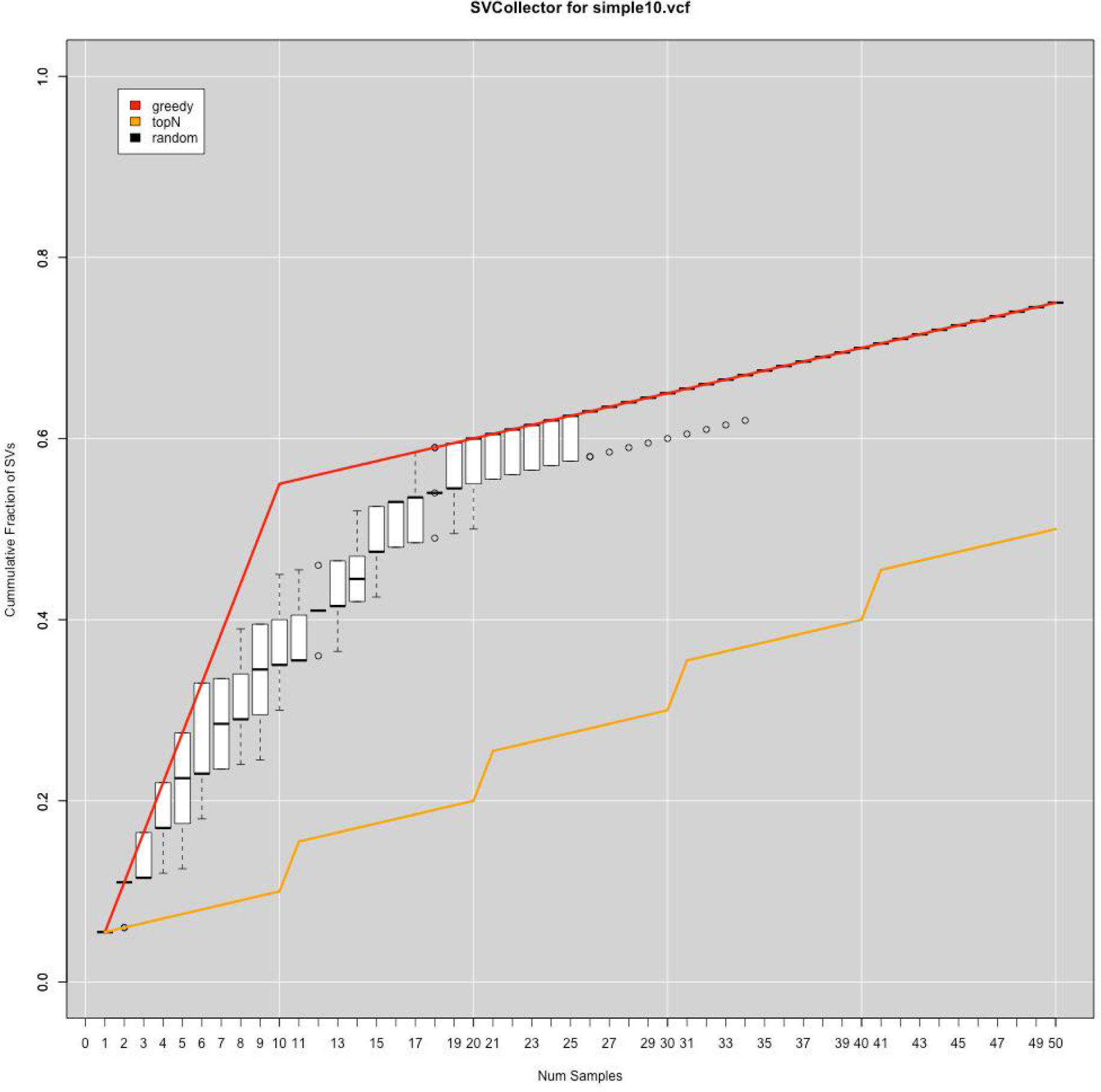
SVCollector curve of a simple simulated population with most variants shared within the population group.

**Supplemental Figure 2:**
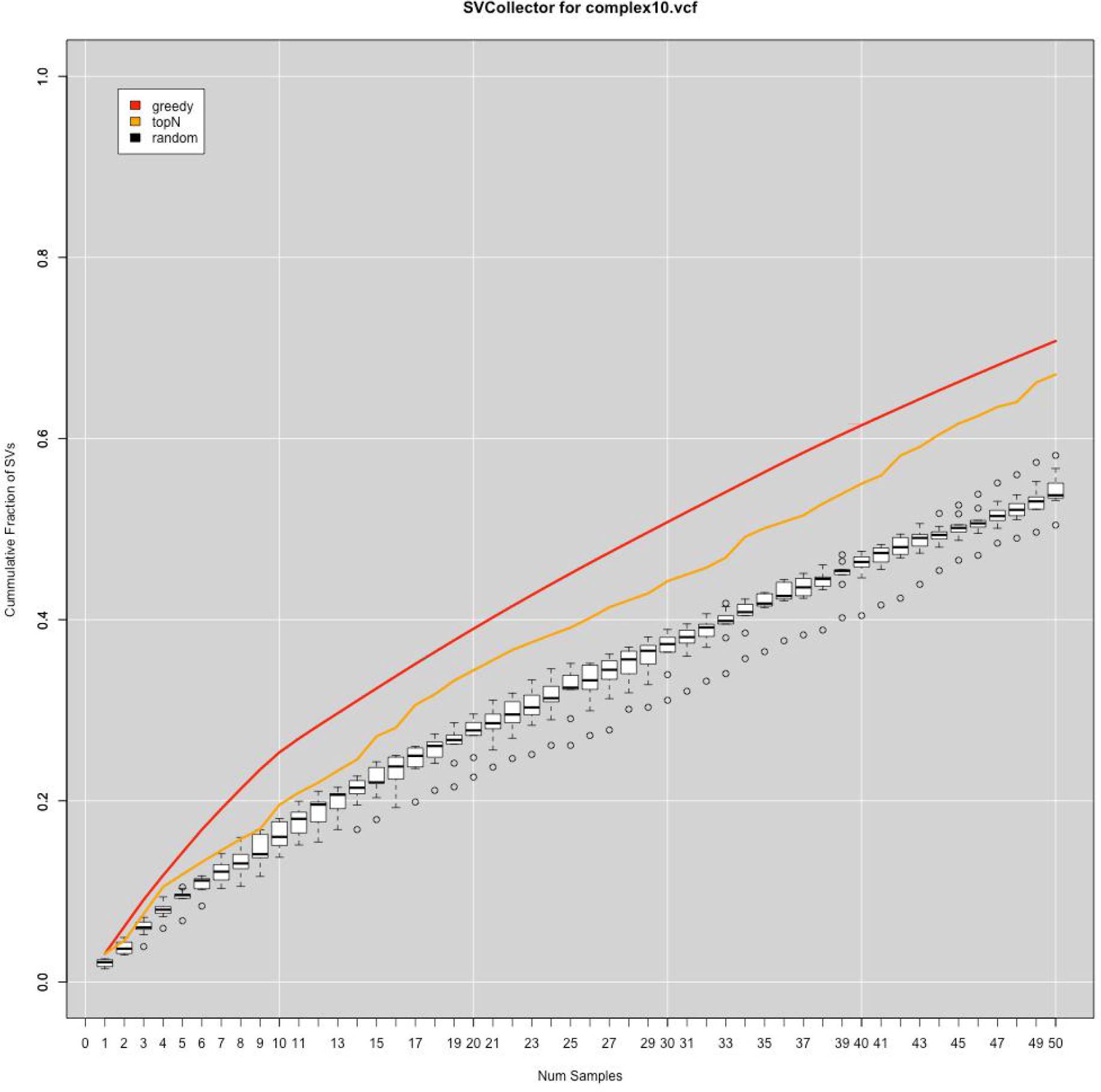
SVCollector curve of a complex simulated population with most variants specific to the individuals.

## Supplemental Note 2. 1000 Genomes Analysis

We downloaded the VCF file from the 1000 genomes FTP site: ftp://ftp.1000genomes.ebi.ac.uk/vol1/ ftp/phase3/integrated_sv_map/ALL.wgs.mergedSV.v8.20130502.svs.genotypes.vcf.gz.

To further assess the robustness of our ranking using the greedy approach we sub-selected once 70% (46,589 SVs) of the SVs reported over the 1000 genomes and once 50% (33,278 SVs) of the SVs. Next, we compared the selection of these sub-selected lists of SVs to the original ranking based on all SVs. For the 70% subsampled set we observed that 7 out of the top 10 accessions are the same compared to the full text. This ratio remains similar for the top 100 ranked samples out of which 71 samples where the same. For the 50% subsampled VCF file we observed similar numbers where 7 out of 10 remained the same within the top 10 ranked samples. For the top 100 this ratio dropped to 59 out of the 100 samples. This illustrates the robustness of SVCollector when it comes to sensitivity issues for SV calling as the ranking remains very similar even with only 50% of the SVs present.

**Supplemental Figure 3:**
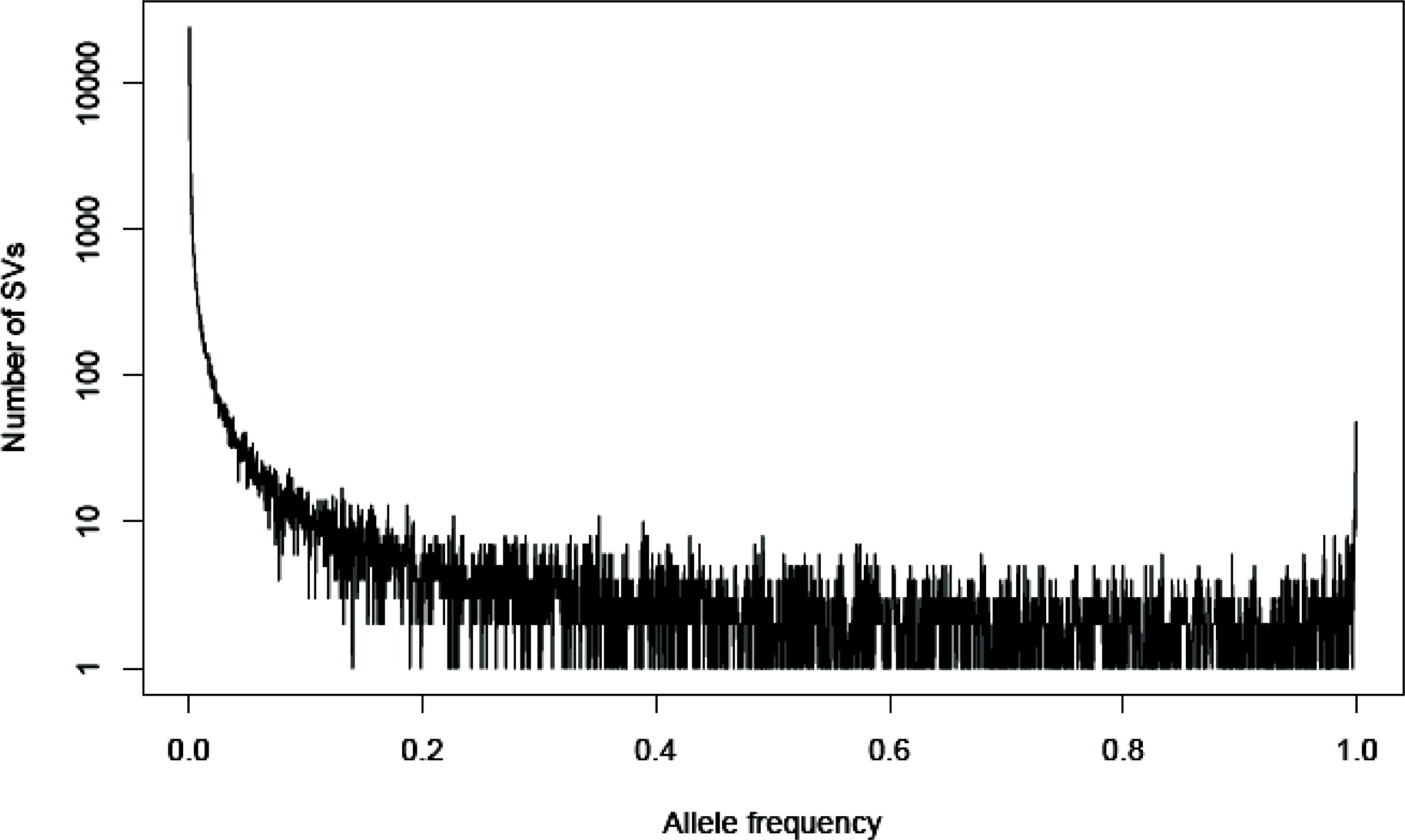
Allele frequency distribution across the 1000 Genomes samples

**Supplemental Figure 4:**
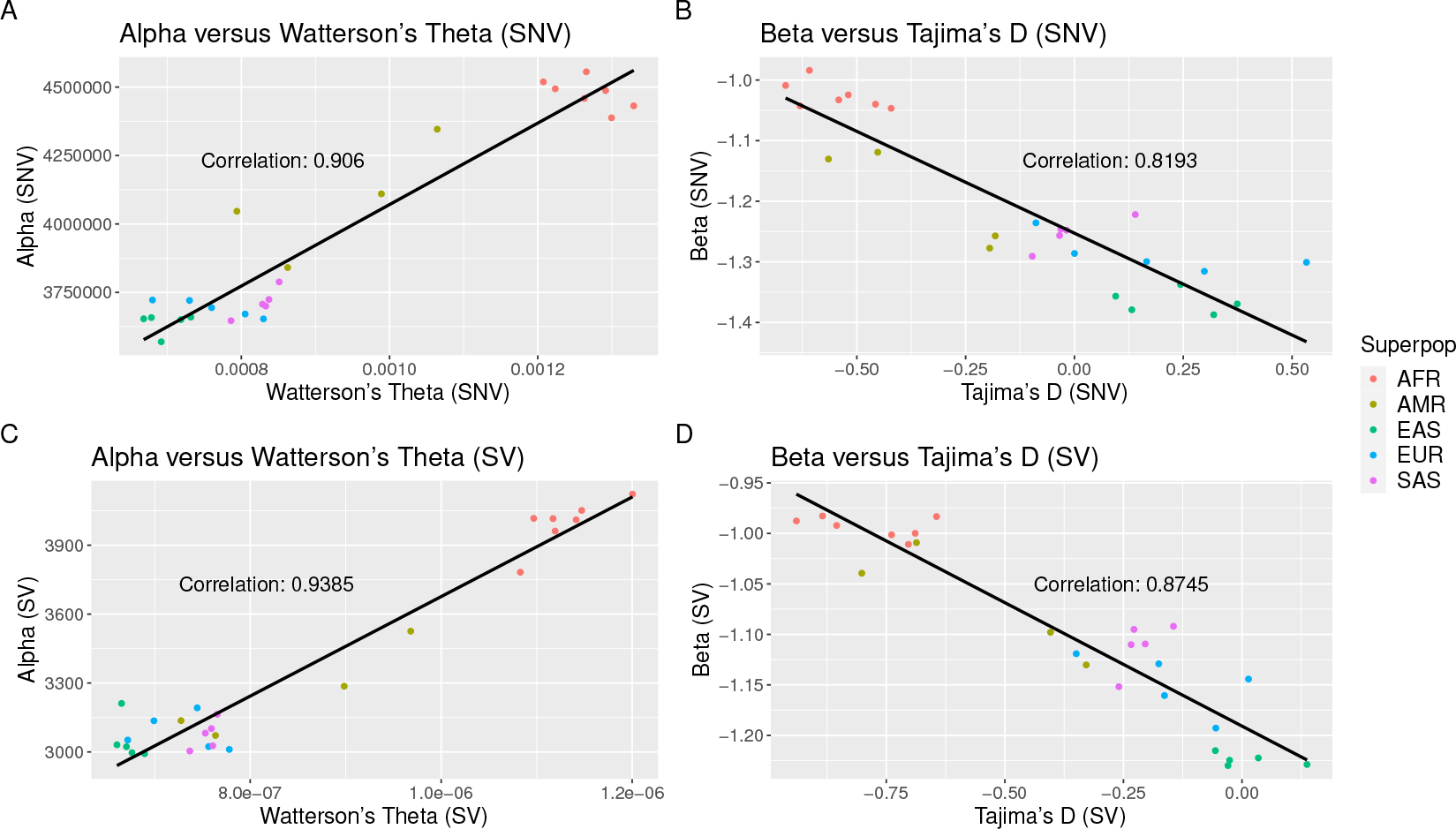
Linear regressions of new versus established population diversity metrics. Each point represents one of the 26 subpopulations in the 1000 Genomes Project. The upper panels are calculated on SNV data and the lower panels are calculated on SV data.

**Supplemental Figure 5:**
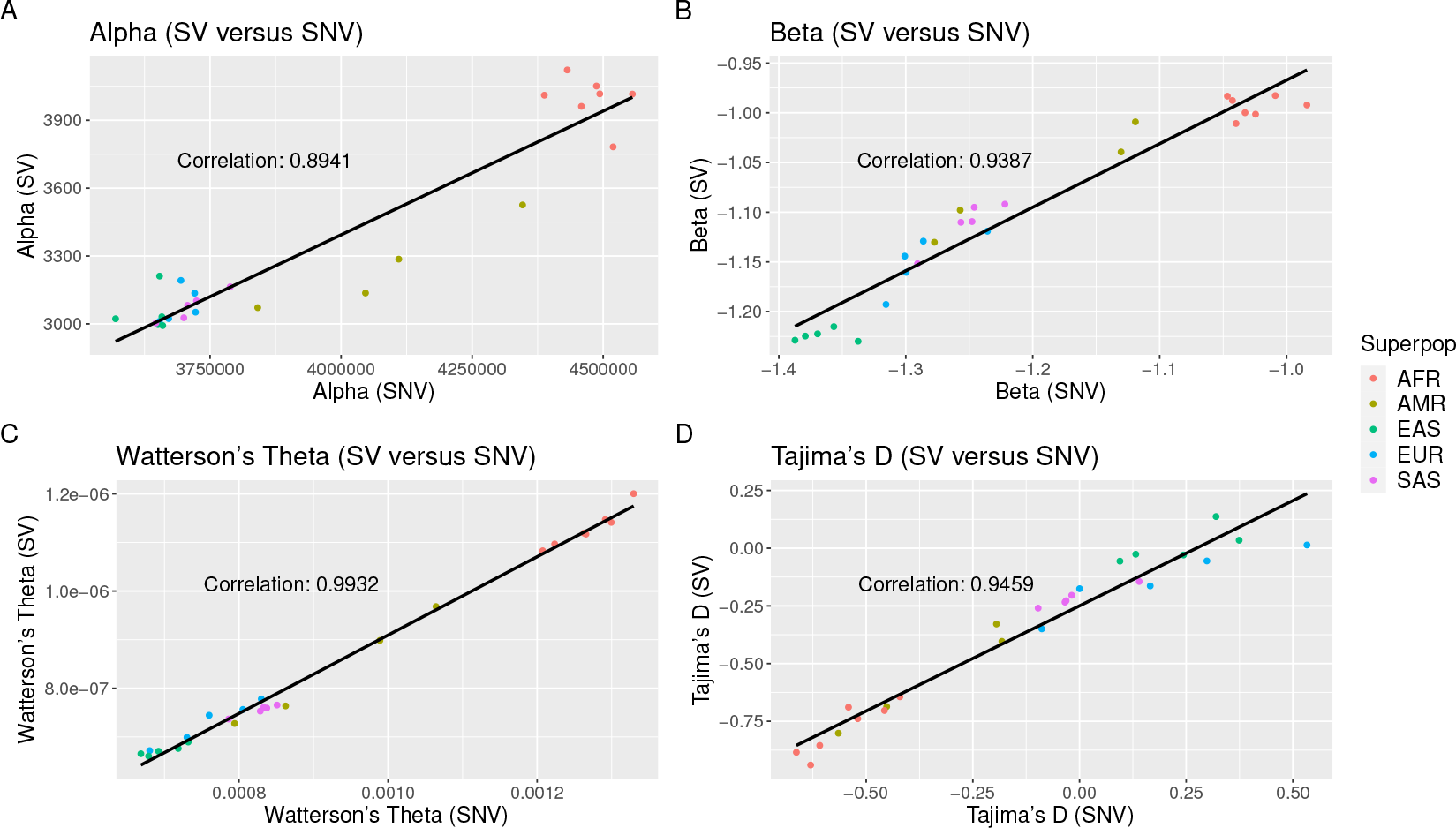
Linear regressions of population diversity metrics (SV versus SNV) for each of the 26 subpopulations.

**Supplemental Figure 6:**
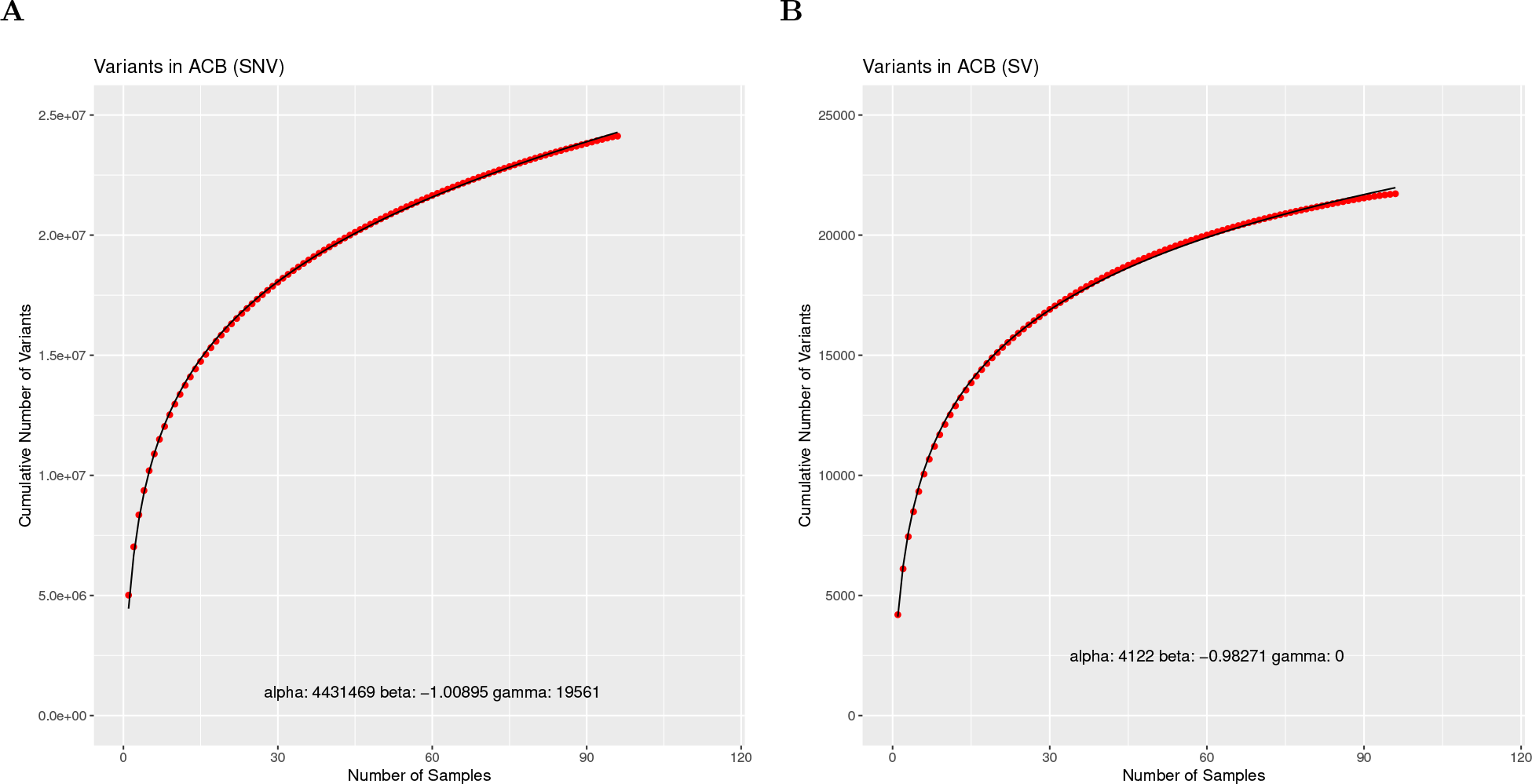
SVCollector curves for the African Caribbeans in Barbados (ACB) population using **(A)** SNV data and **(B)** SV data.

**Supplemental Figure 7:**
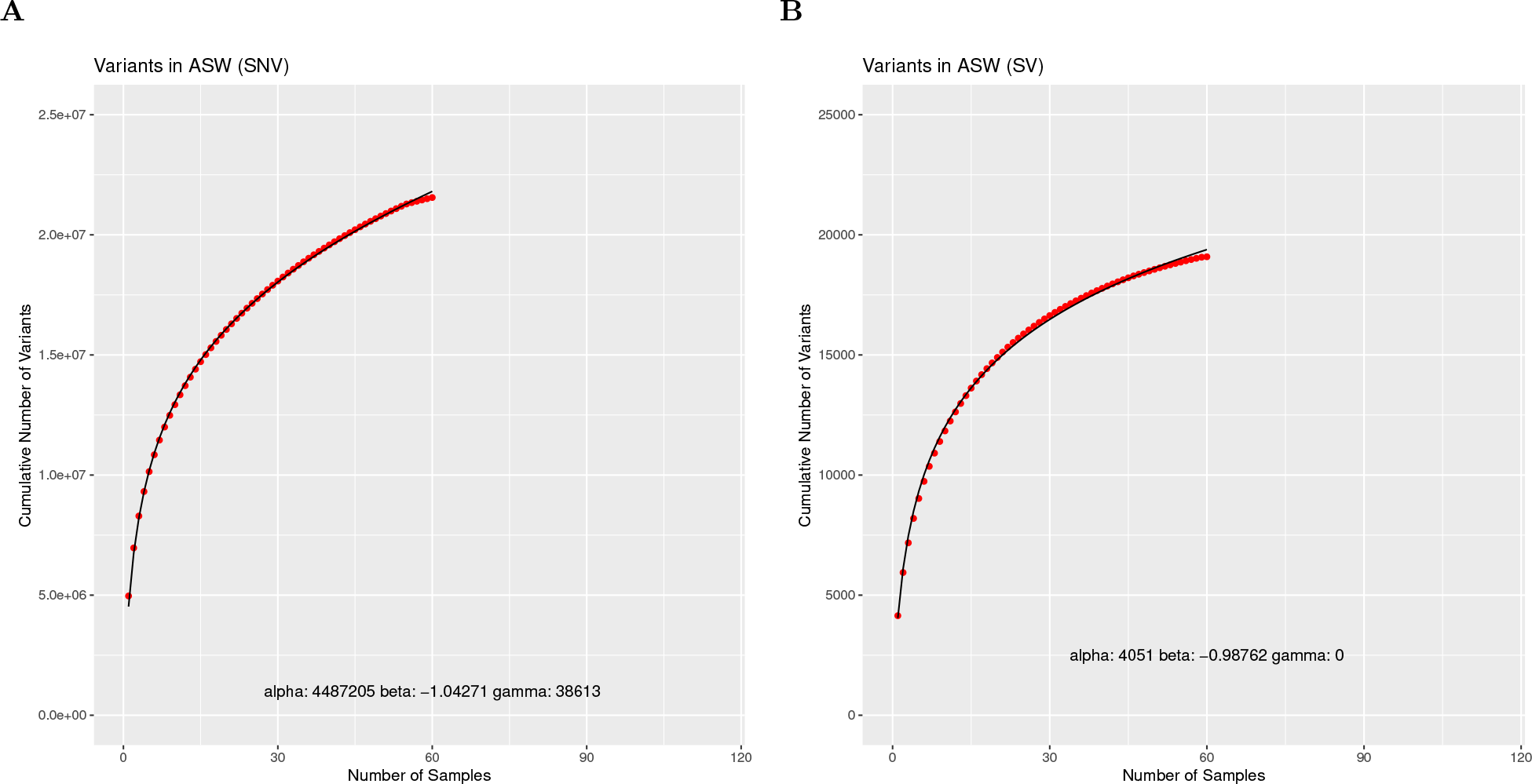
SVCollector curves for the Americans of African Ancestry in SW USA (ASW) population using **(A)** SNV data and **(B)** SV data.

**Supplemental Figure 8:**
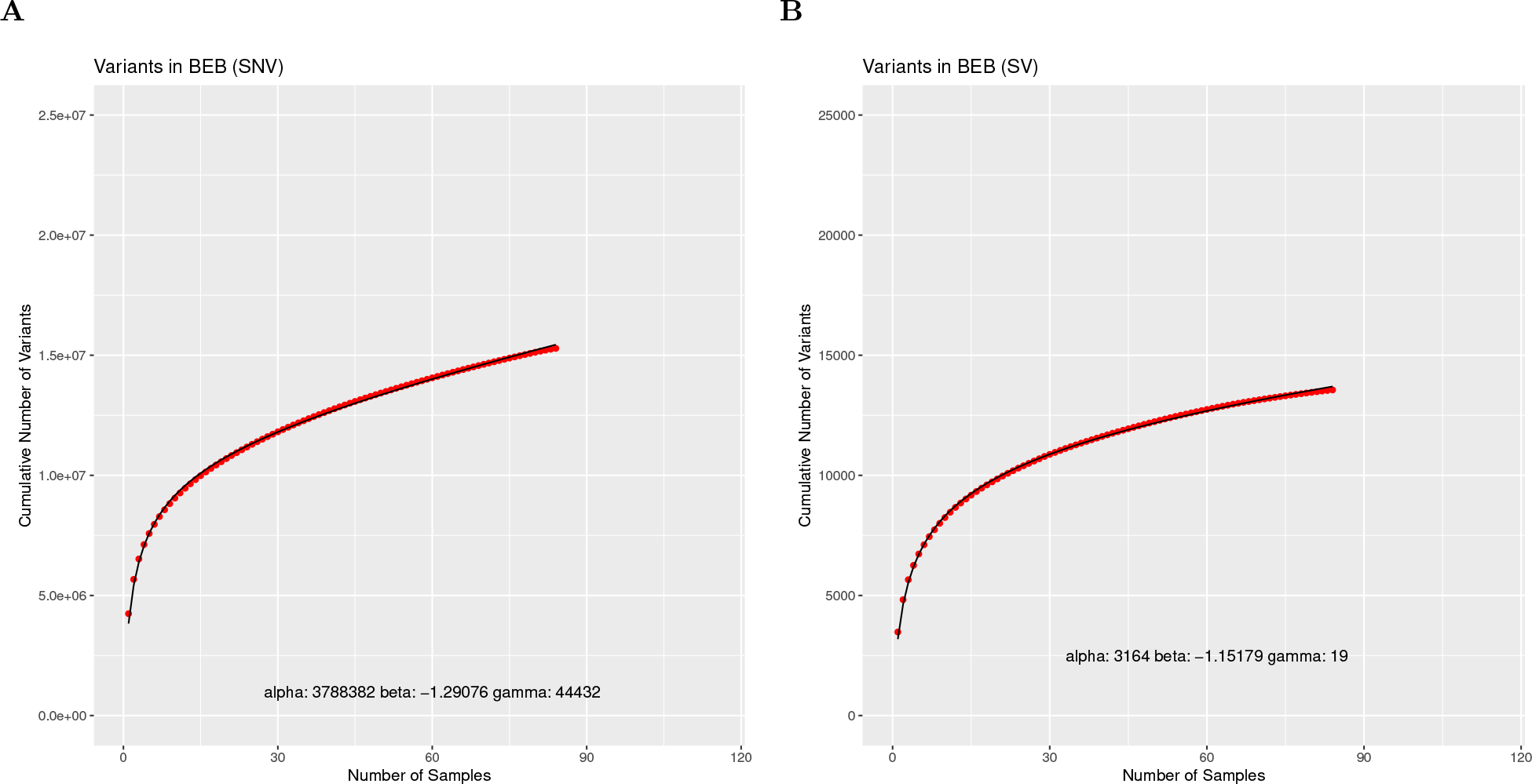
SVCollector curves for the Bengali from Bangladesh (BEB) population using **(A)** SNV data and **(B)** SV data.

**Supplemental Figure 9:**
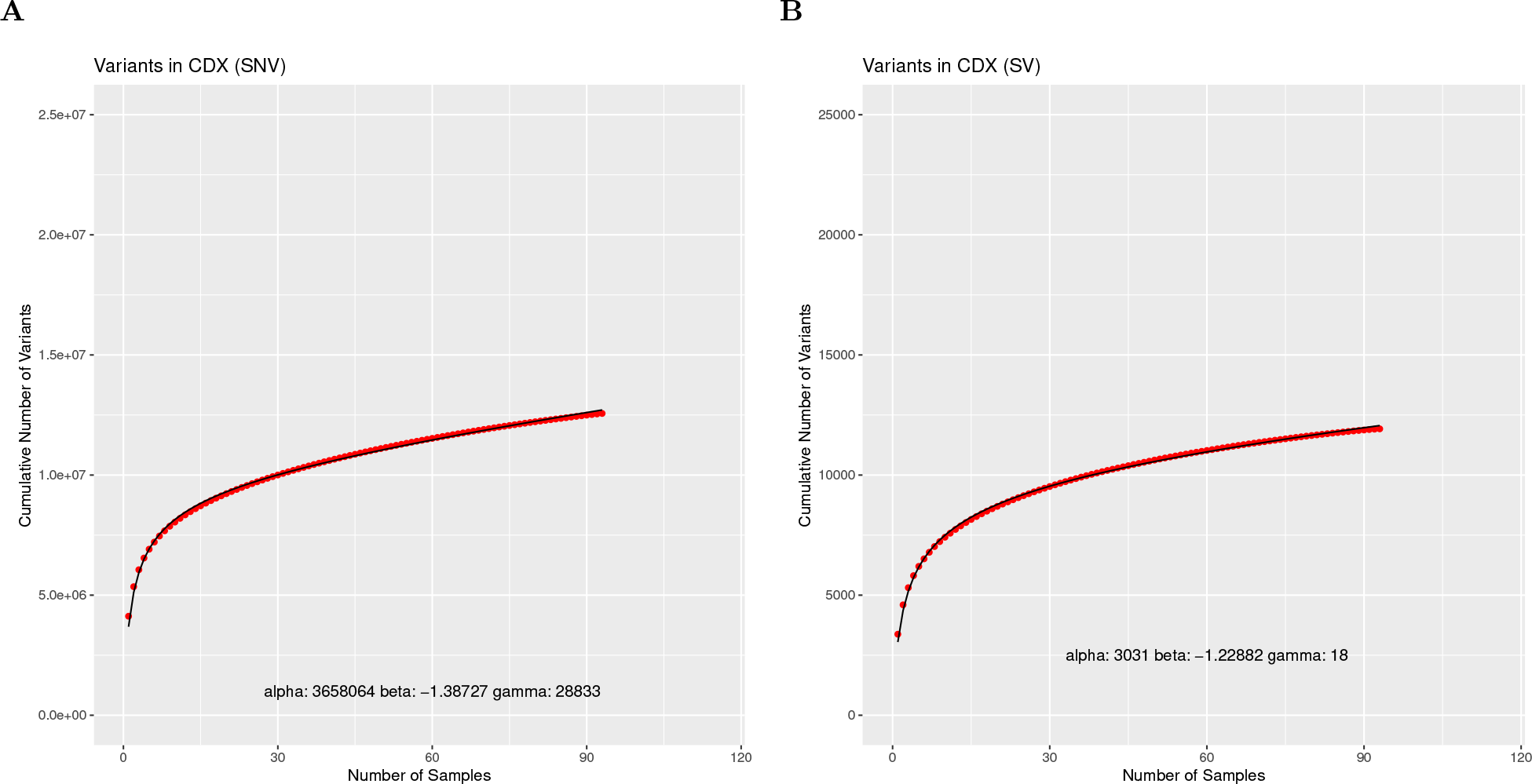
SVCollector curves for the Chinese Dai in Xishuangbanna, China (CDX) population using **(A)** SNV data and **(B)** SV data.

**Supplemental Figure 10:**
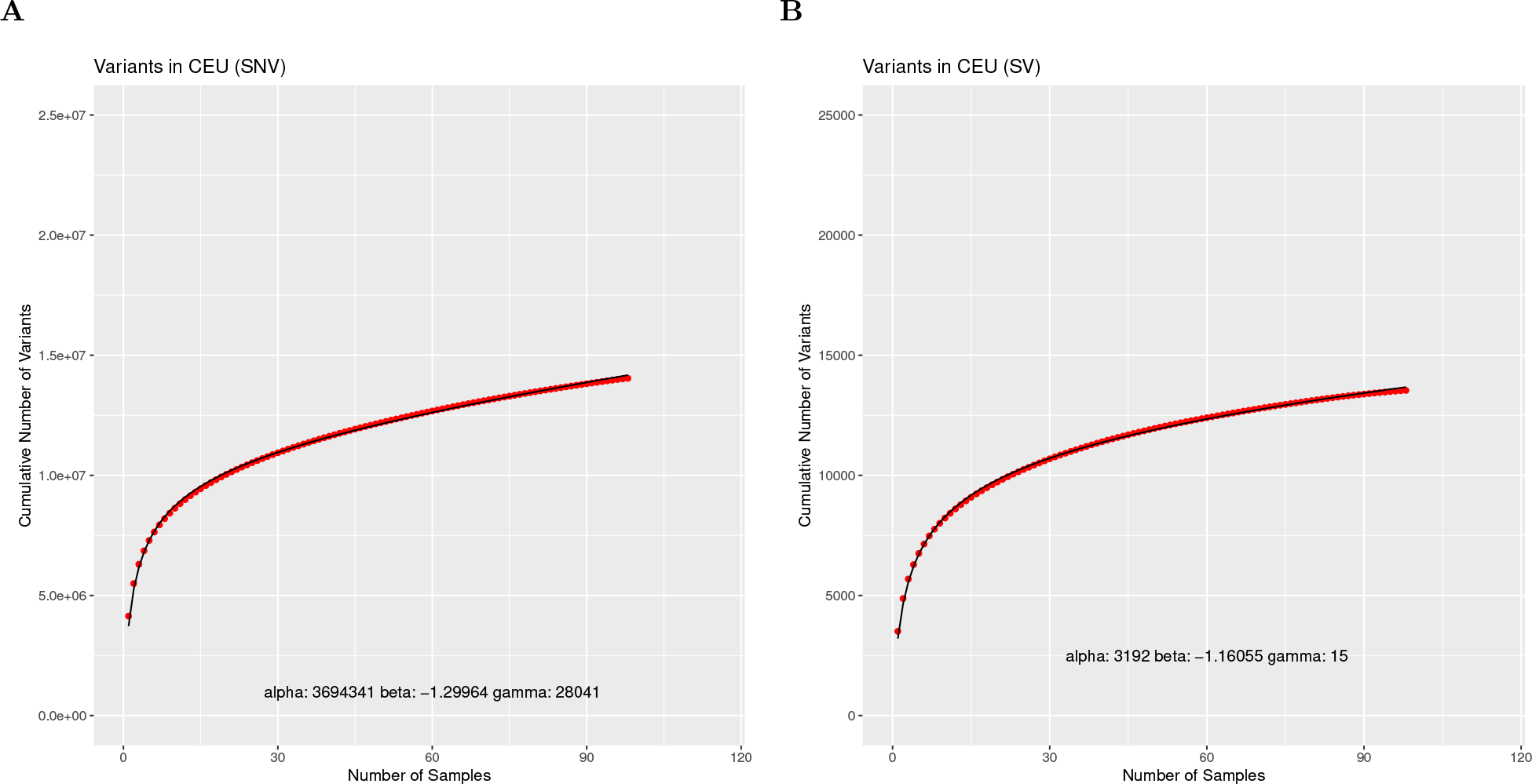
SVCollector curves for the Utah Residents (CEPH) with Northern and Western European Ancestry (CEU) population using **(A)** SNV data and **(B)** SV data.

**Supplemental Figure 11:**
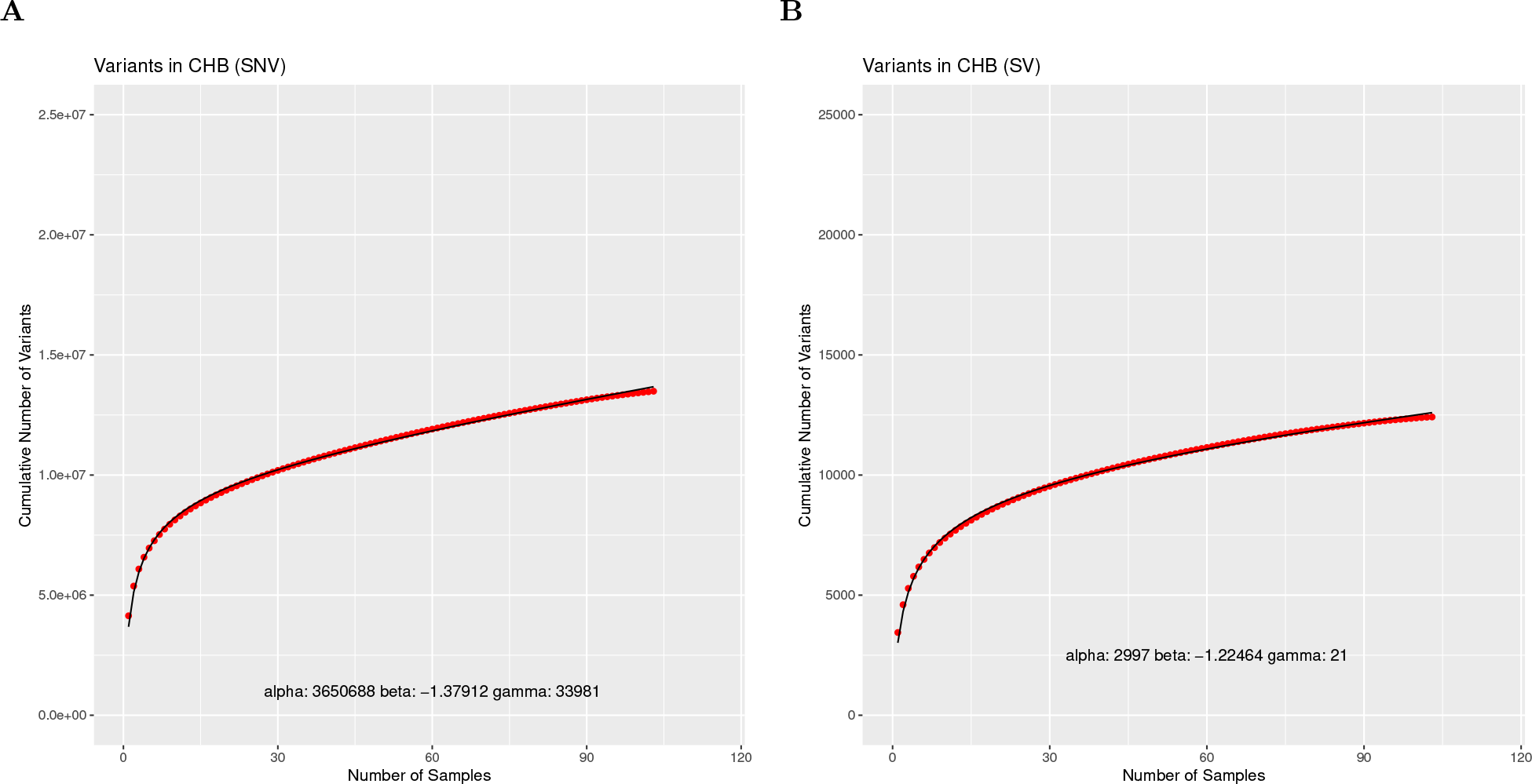
SVCollector curves for the Han Chinese in Beijing, China (CHB) population using **(A)** SNV data and **(B)** SV data.

**Supplemental Figure 12:**
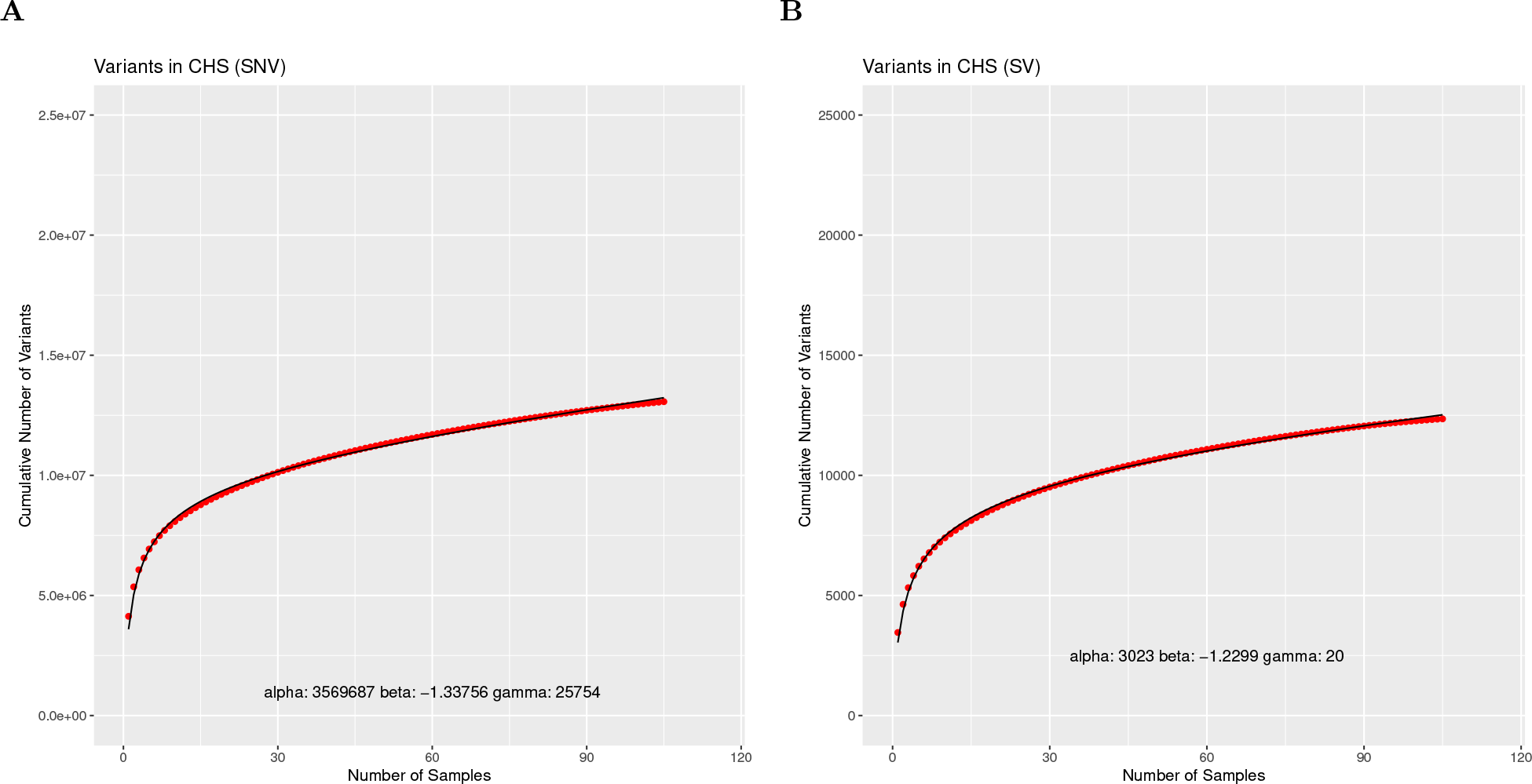
SVCollector curves for the Southern Han Chinese (CHS) population using **(A)** SNV data and SV data.

**Supplemental Figure 13:**
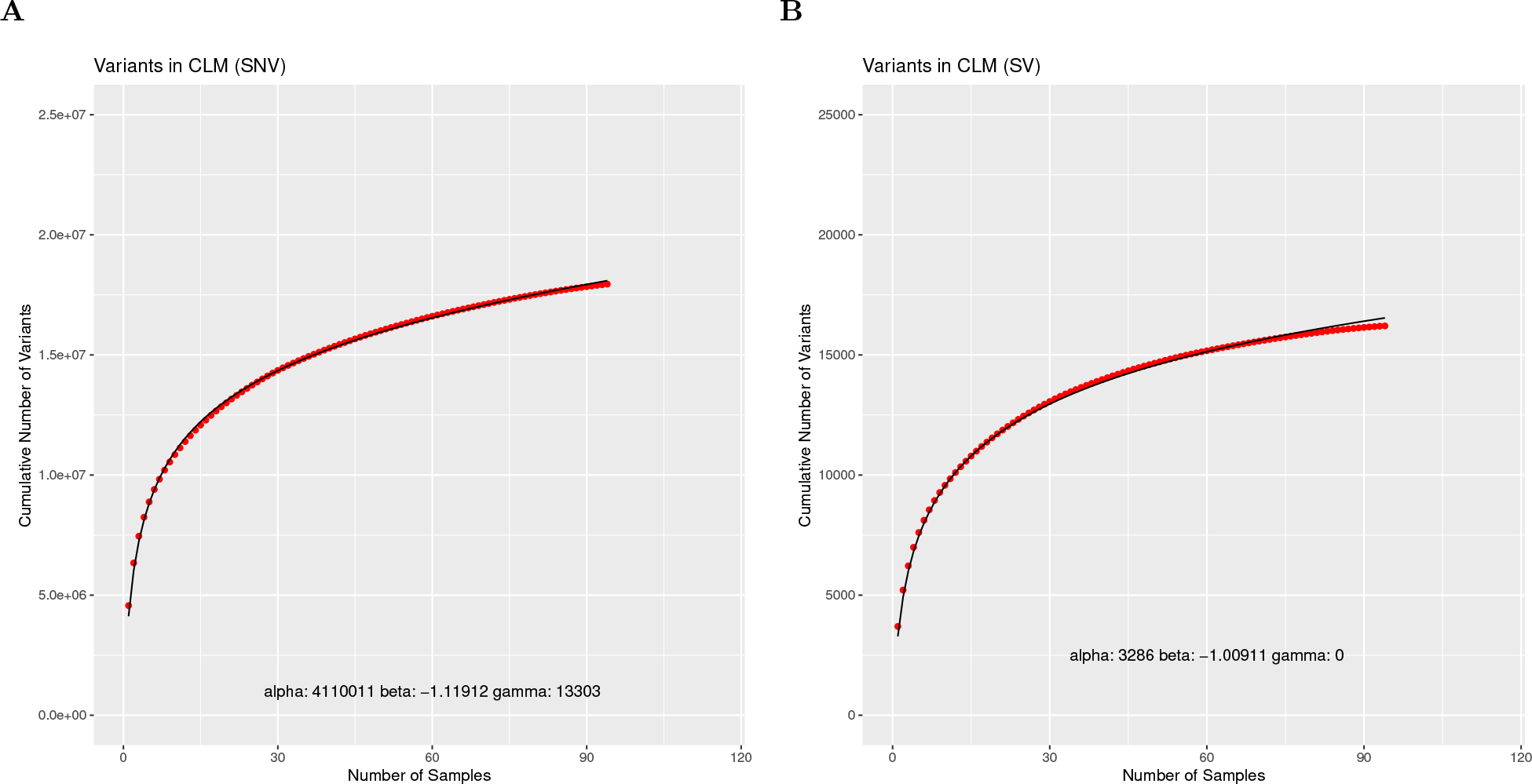
SVCollector curves for the Colombians from Medellin, Colombia (CLM) population using **(A)** SNV data and **(B)** SV data.

**Supplemental Figure 14:**
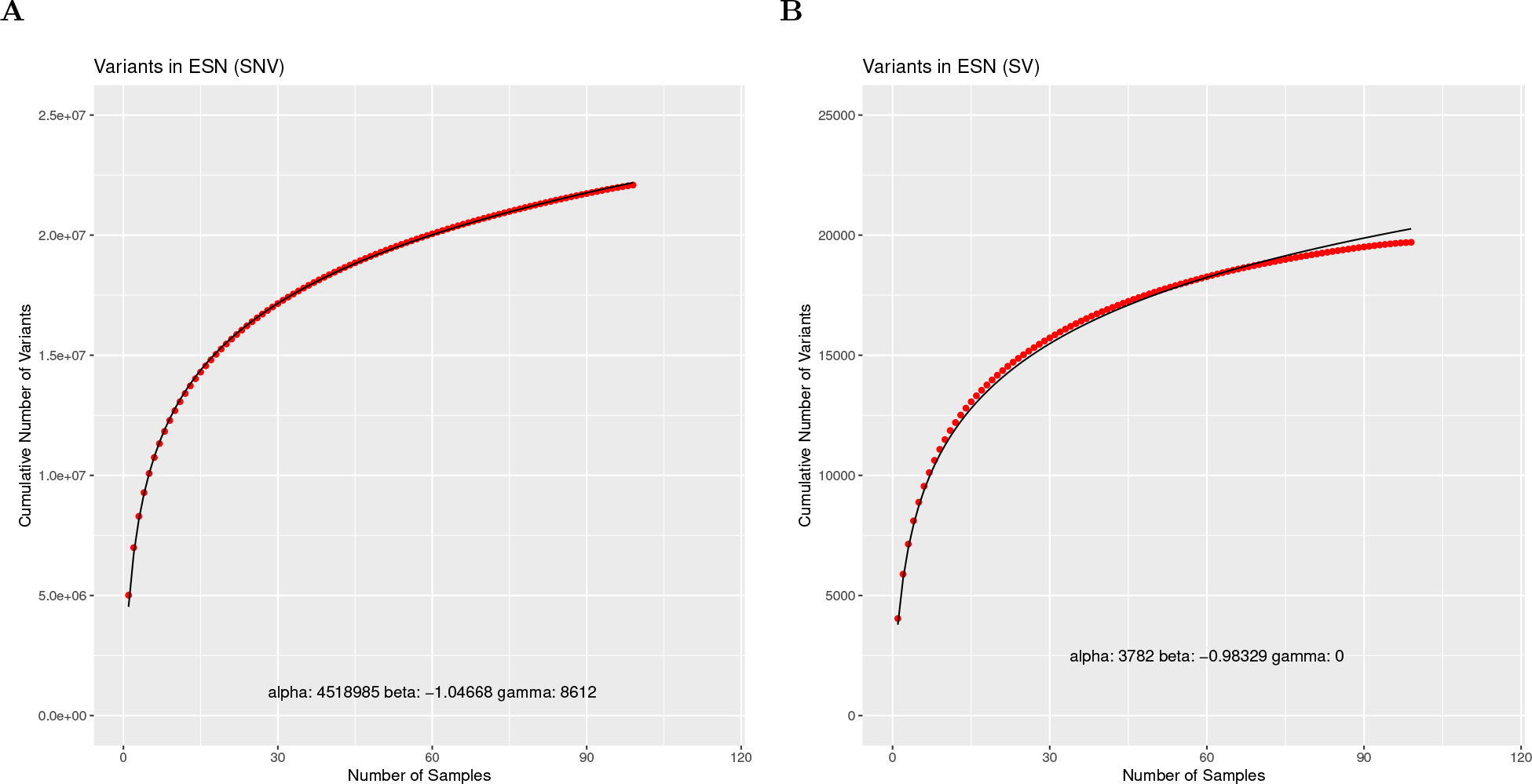
SVCollector curves for the Esan in Nigeria (ESN) population using **(A)** SNV data and **(B)** SV data.

**Supplemental Figure 15:**
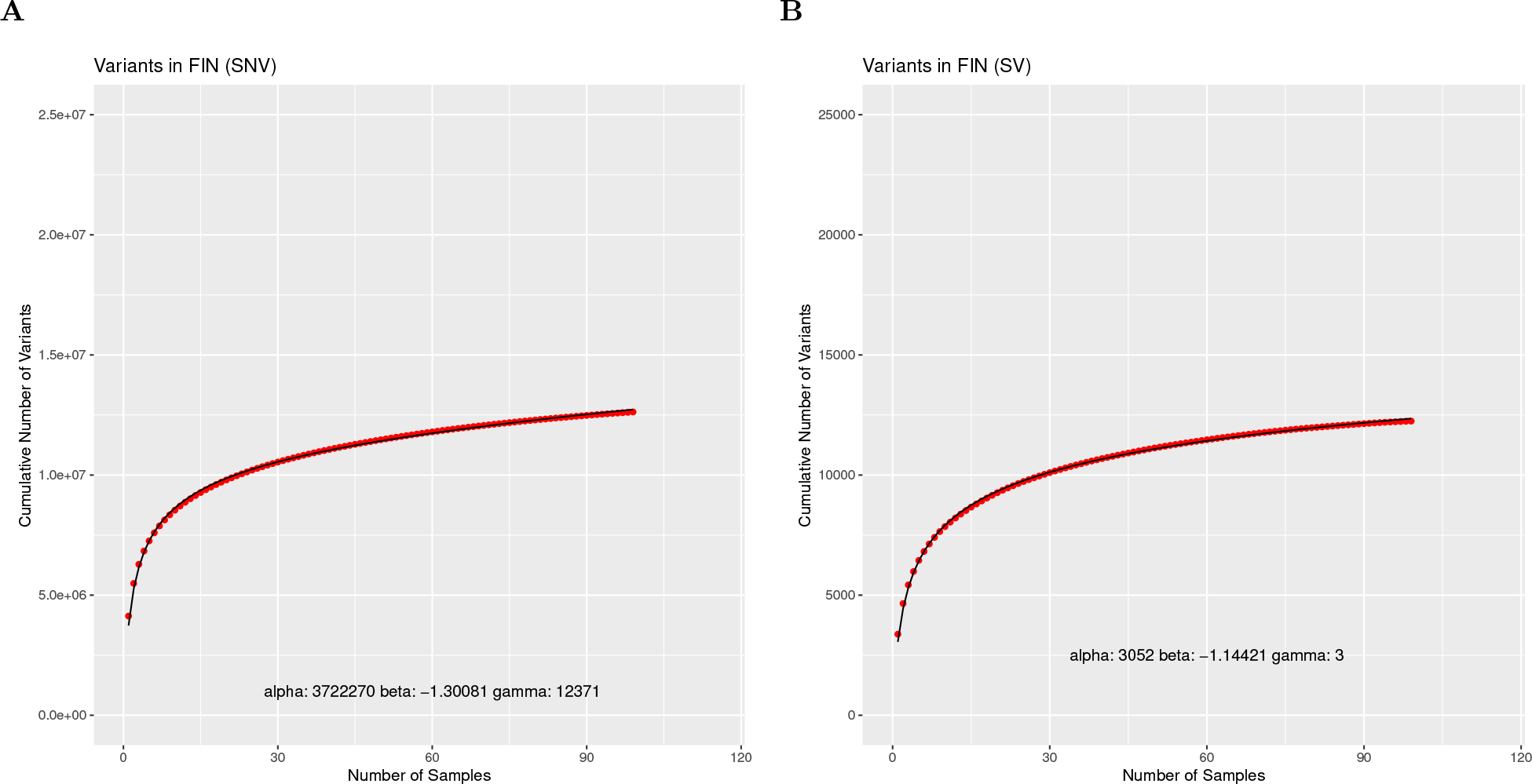
SVCollector curves for the Finnish in Finland (FIN) population using **(A)** SNV data and **(B)** SV data.

**Supplemental Figure 16:**
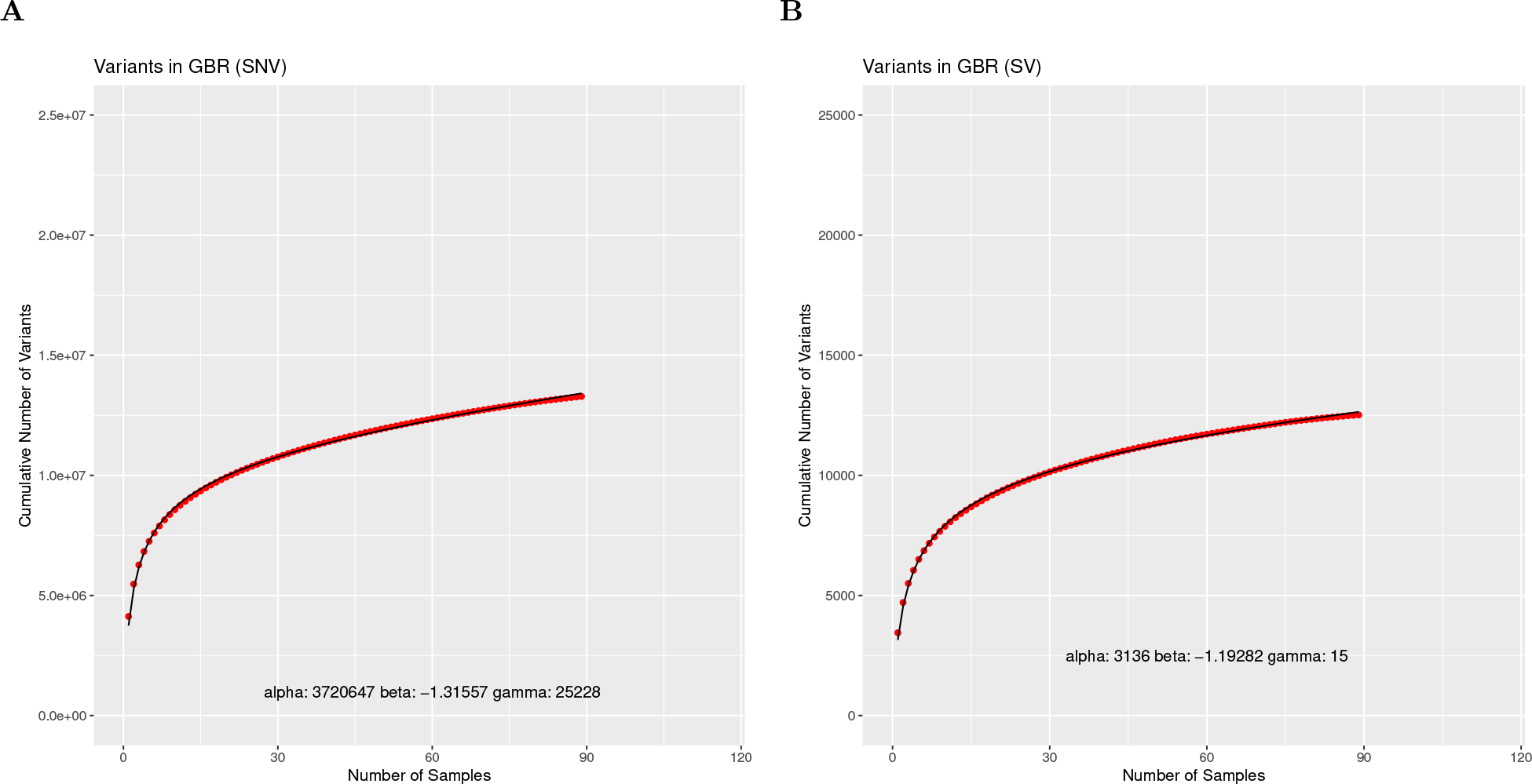
SVCollector curves for the British in England and Scotland (GBR) population using **(A)** SNV data and **(B)** SV data.

**Supplemental Figure 17:**
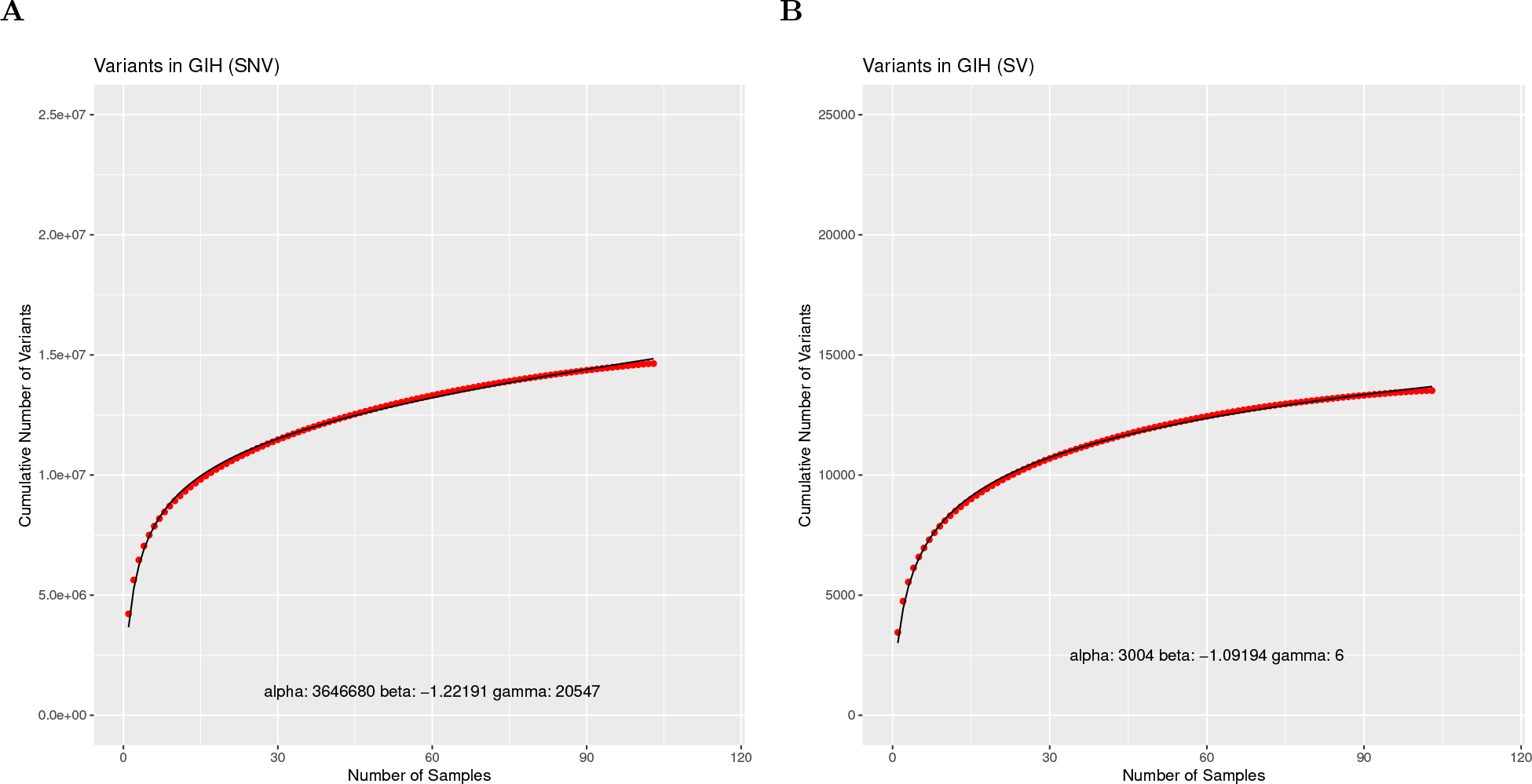
SVCollector curves for the Gujarati Indian from Houston, Texas (GIH) population using **(A)** SNV data and **(B)** SV data.

**Supplemental Figure 18:**
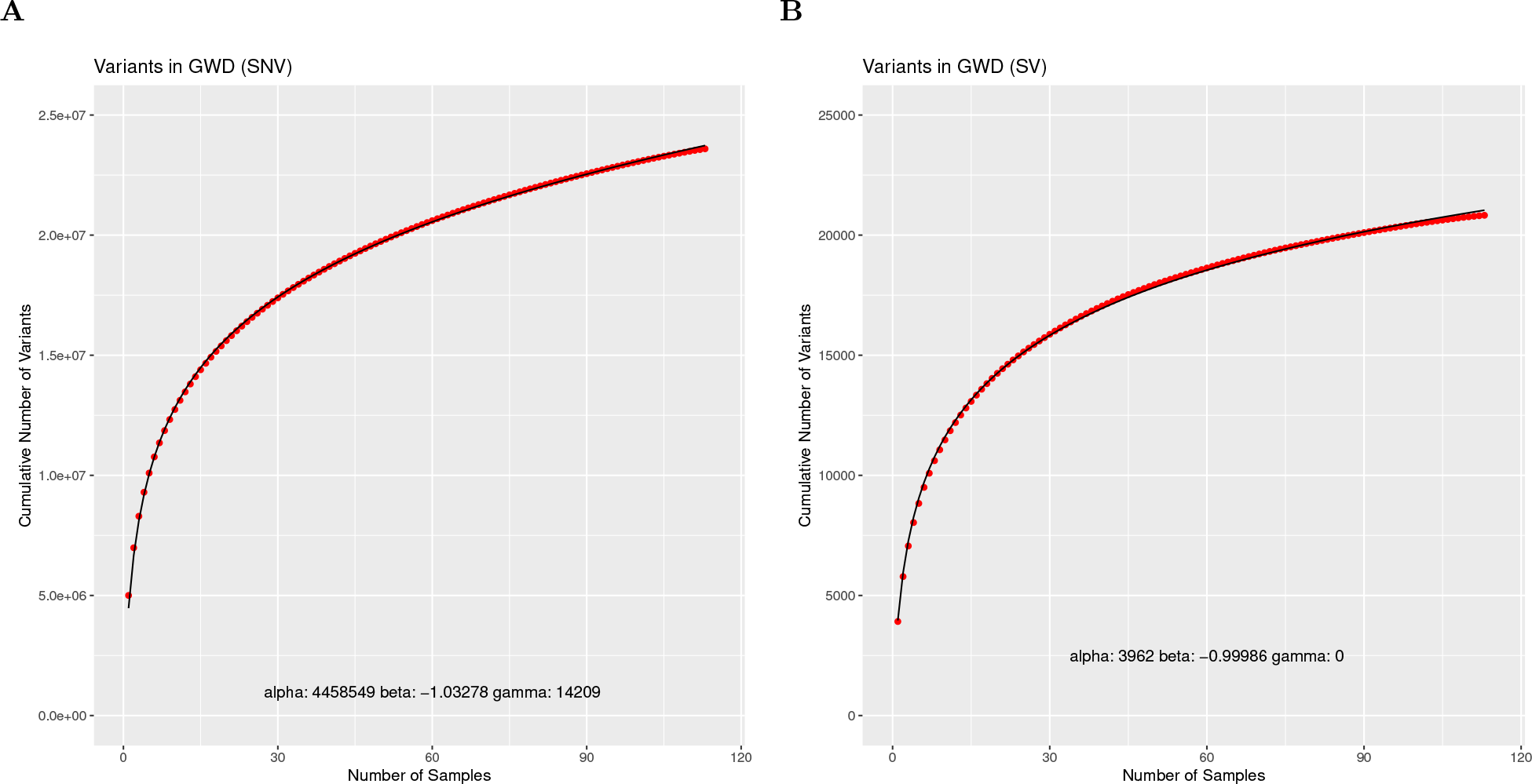
SVCollector curves for the Gambian in Western Divisions in the Gambia (GWD) population using **(A)** SNV data and **(B)** SV data.

**Supplemental Figure 19:**
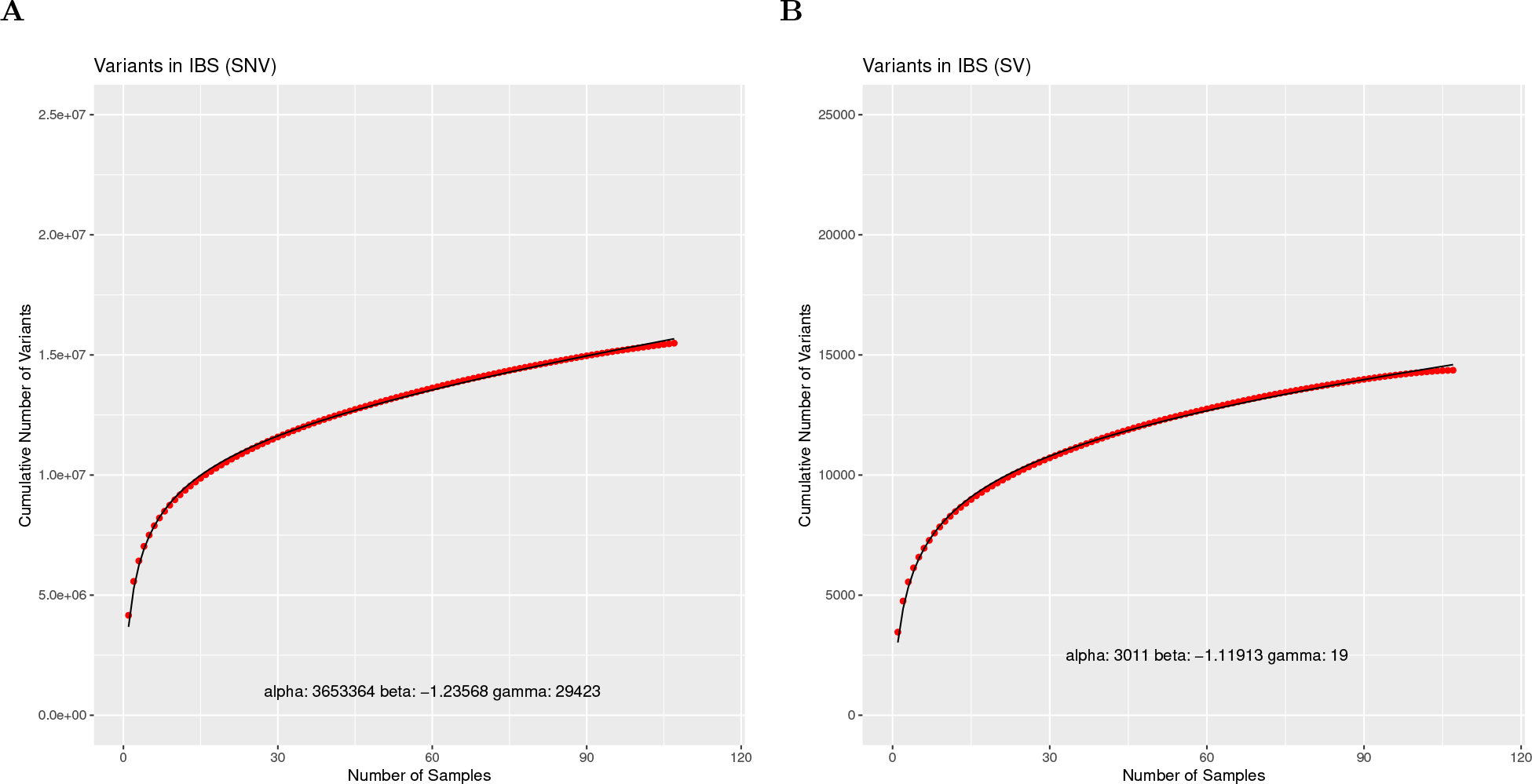
SVCollector curves for the Iberian Population in Spain (IBS) population using **(A)** SNV data and **(B)** SV data.

**Supplemental Figure 20:**
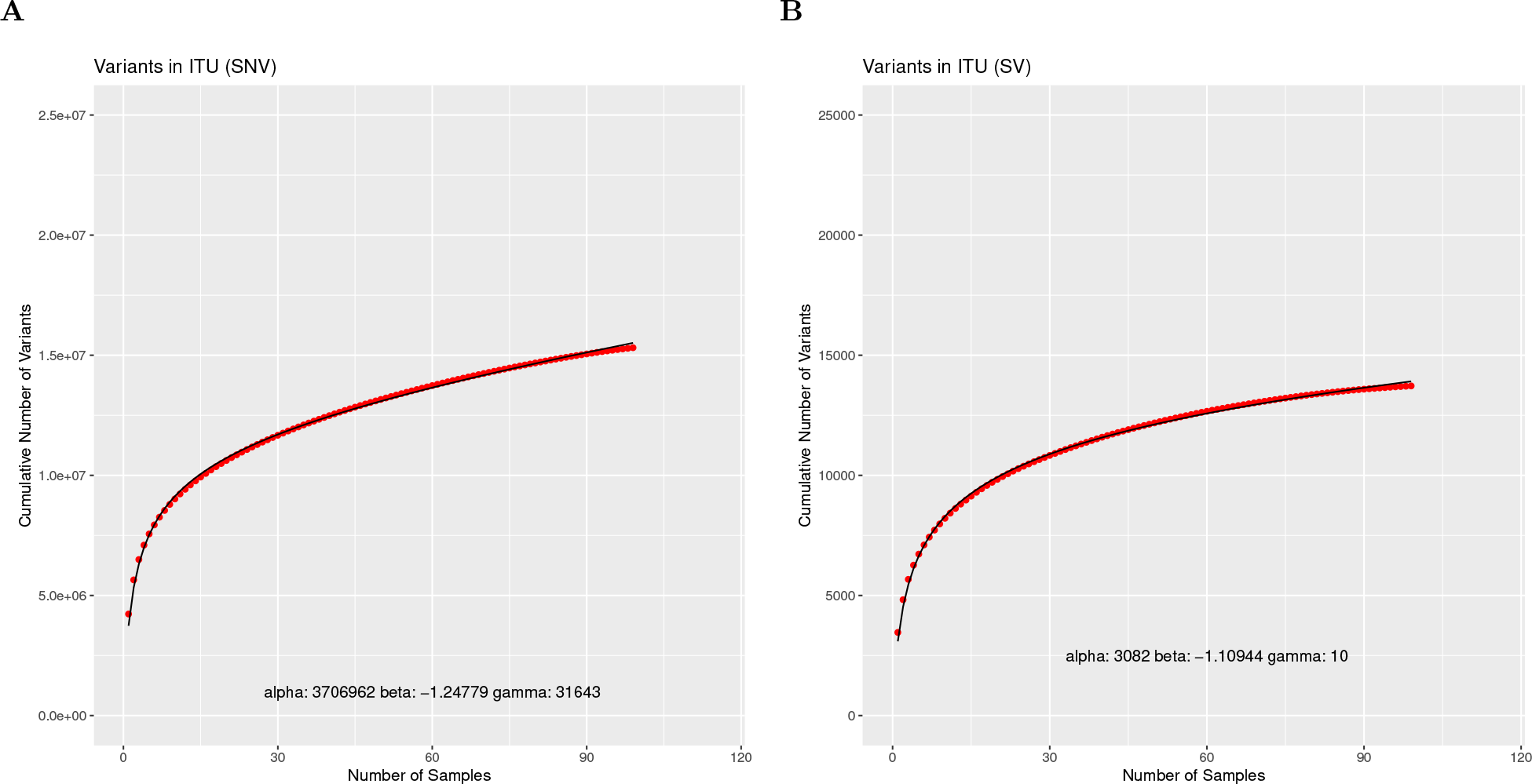
SVCollector curves for the Indian Telugu from the UK (ITU) population using **(A)** SNV data and **(B)** SV data.

**Supplemental Figure 21:**
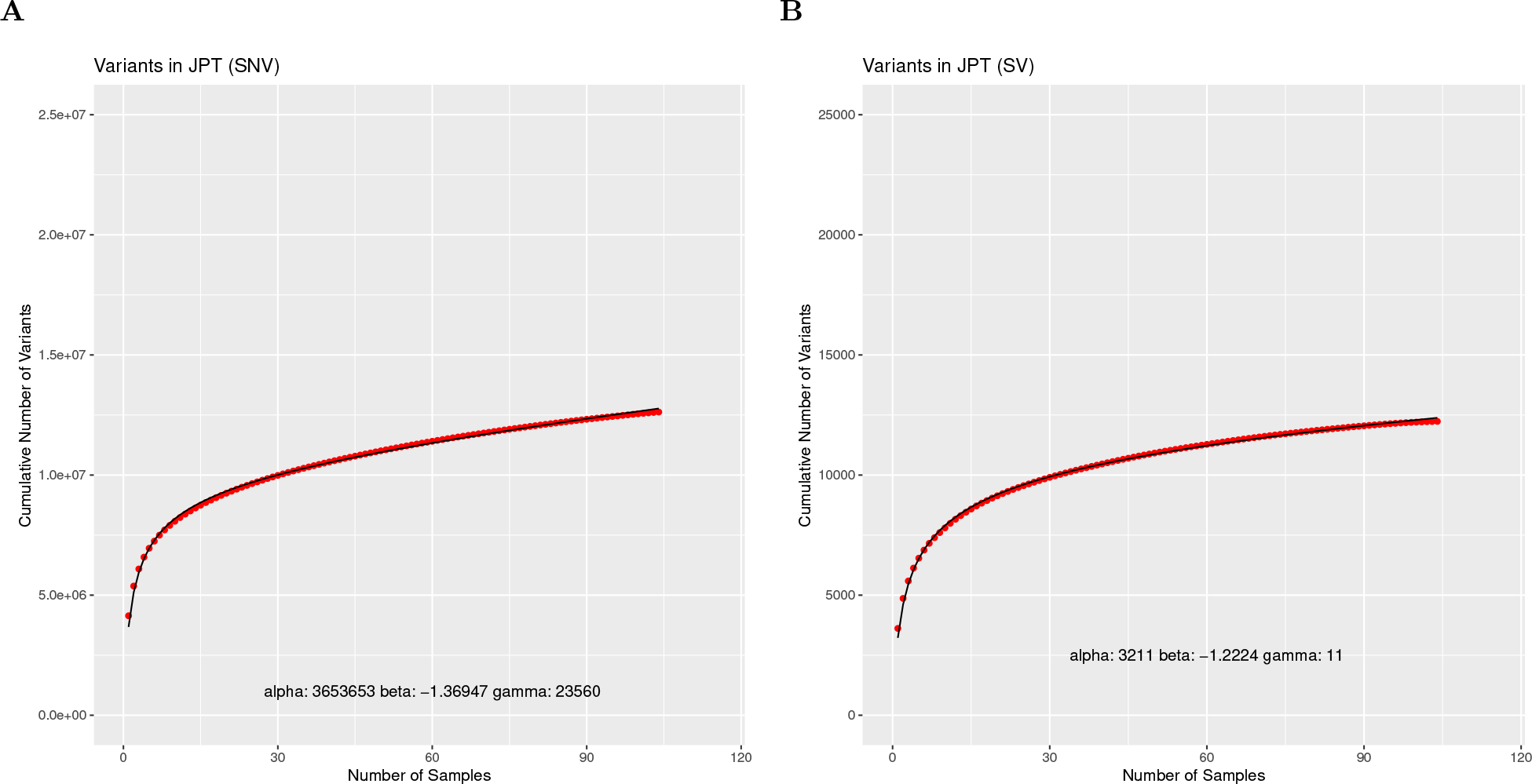
SVCollector curves for the Japanese in Tokyo, Japan (JPT) population using **(A)** SNV data and **(B)** SV data.

**Supplemental Figure 22:**
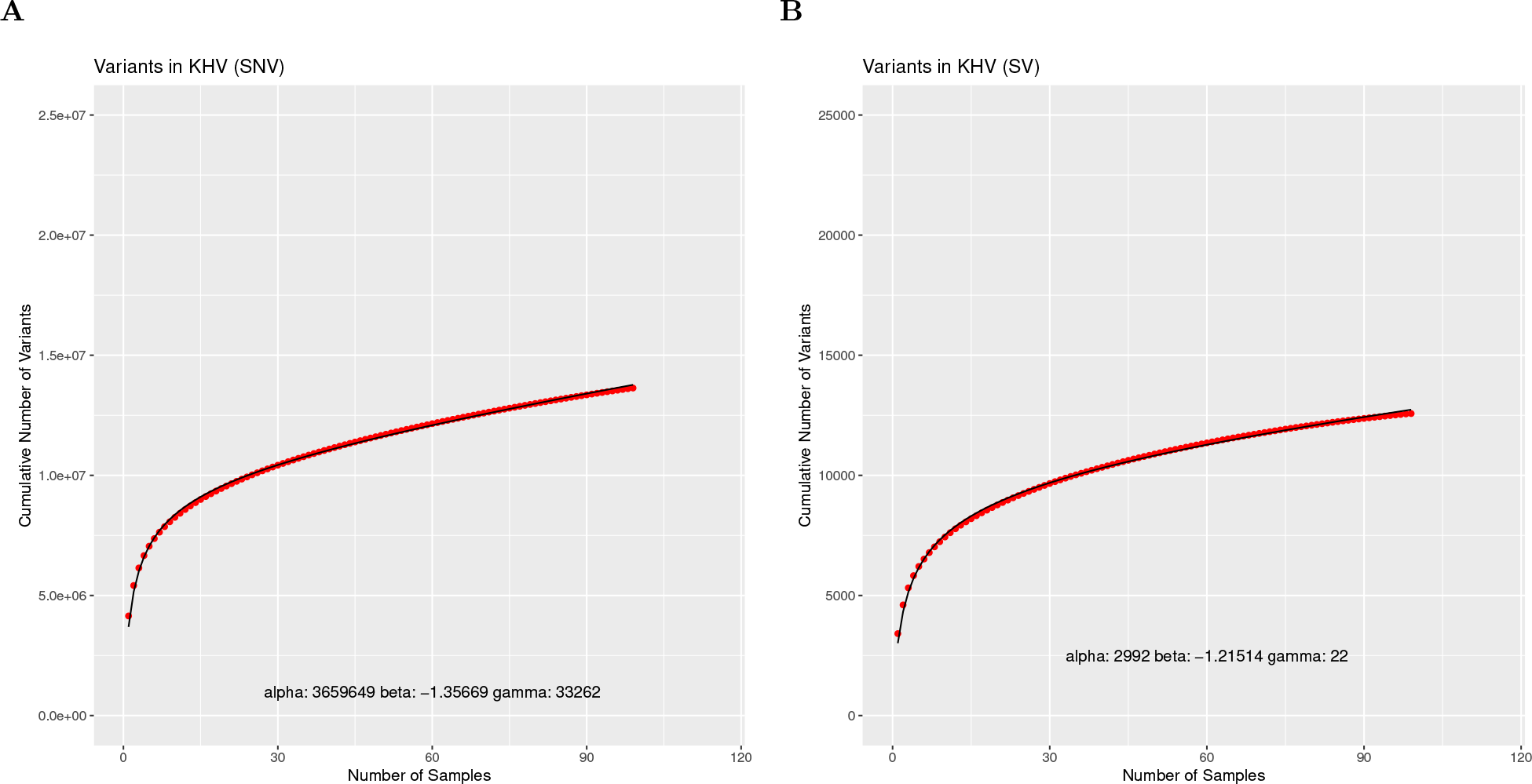
SVCollector curves for the Kinh in Ho Chi Minh City, Vietnam (KHV) population using **(A)** SNV data and **(B)** SV data.

**Supplemental Figure 23:**
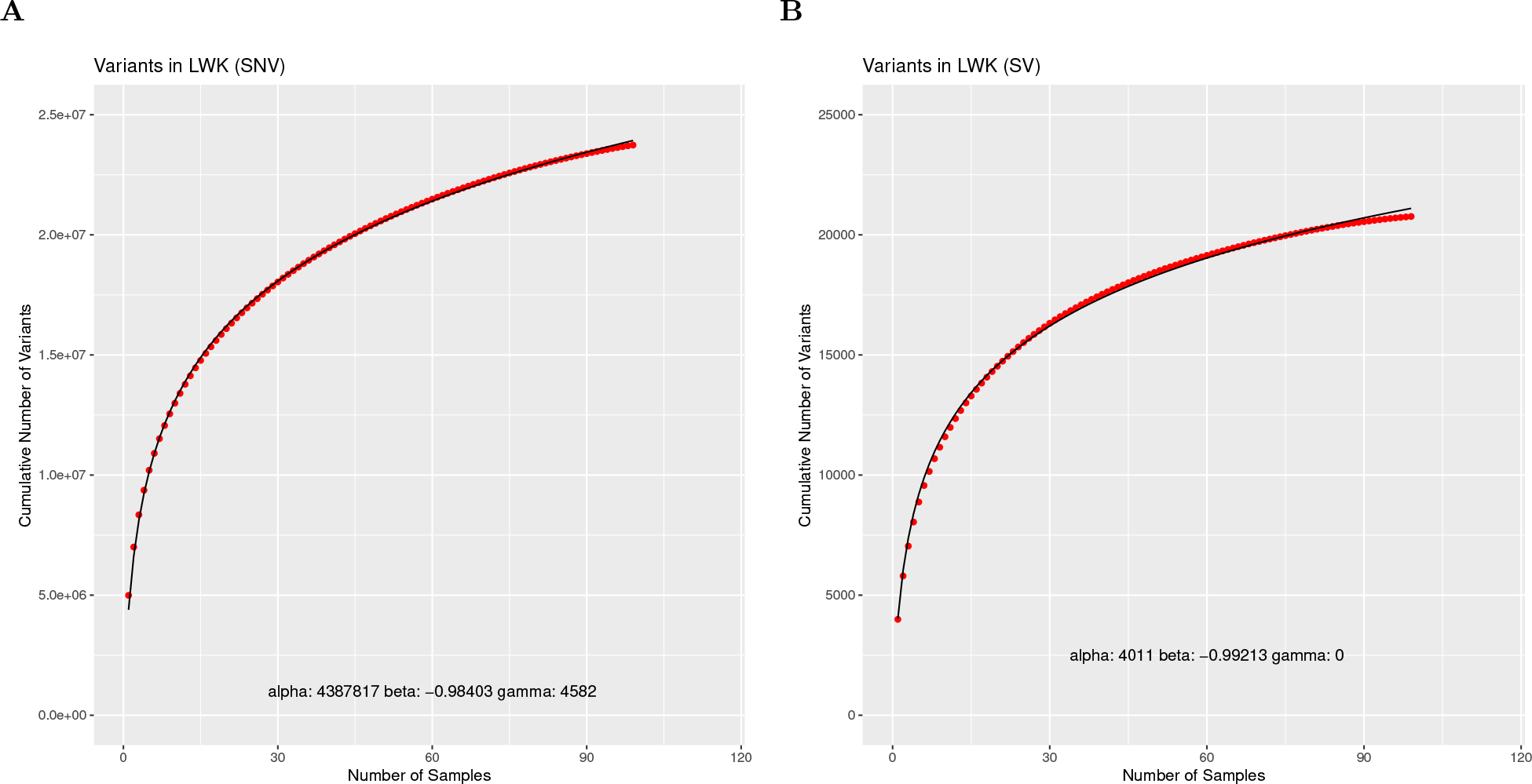
SVCollector curves for the Luhya in Webuye, Kenya (LWK) population using **(A)** SNV data and **(B)** SV data.

**Supplemental Figure 24:**
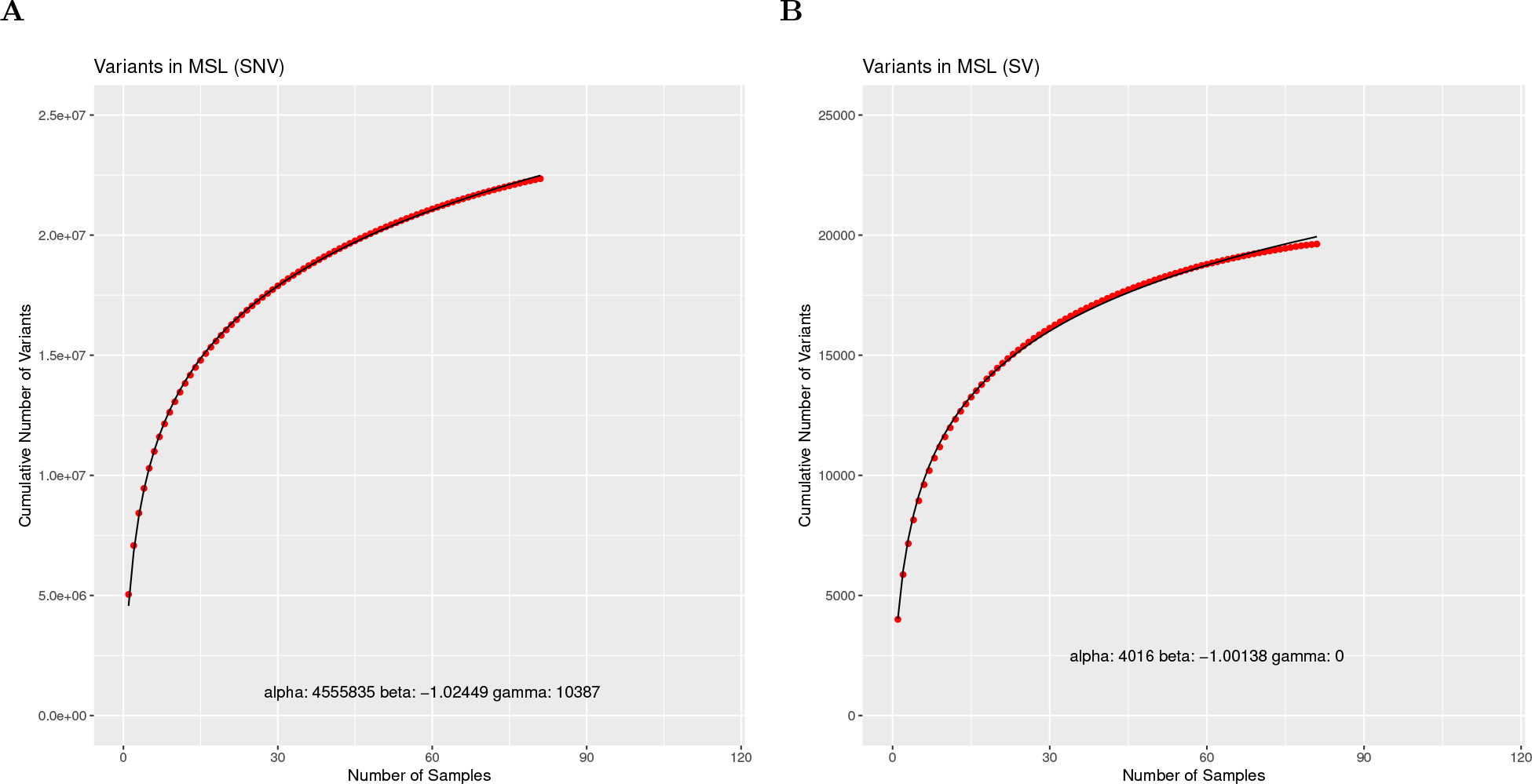
SVCollector curves for the Mende in Sierra Leone (MSL) population using **(A)** SNV data and **(B)** SV data.

**Supplemental Figure 25:**
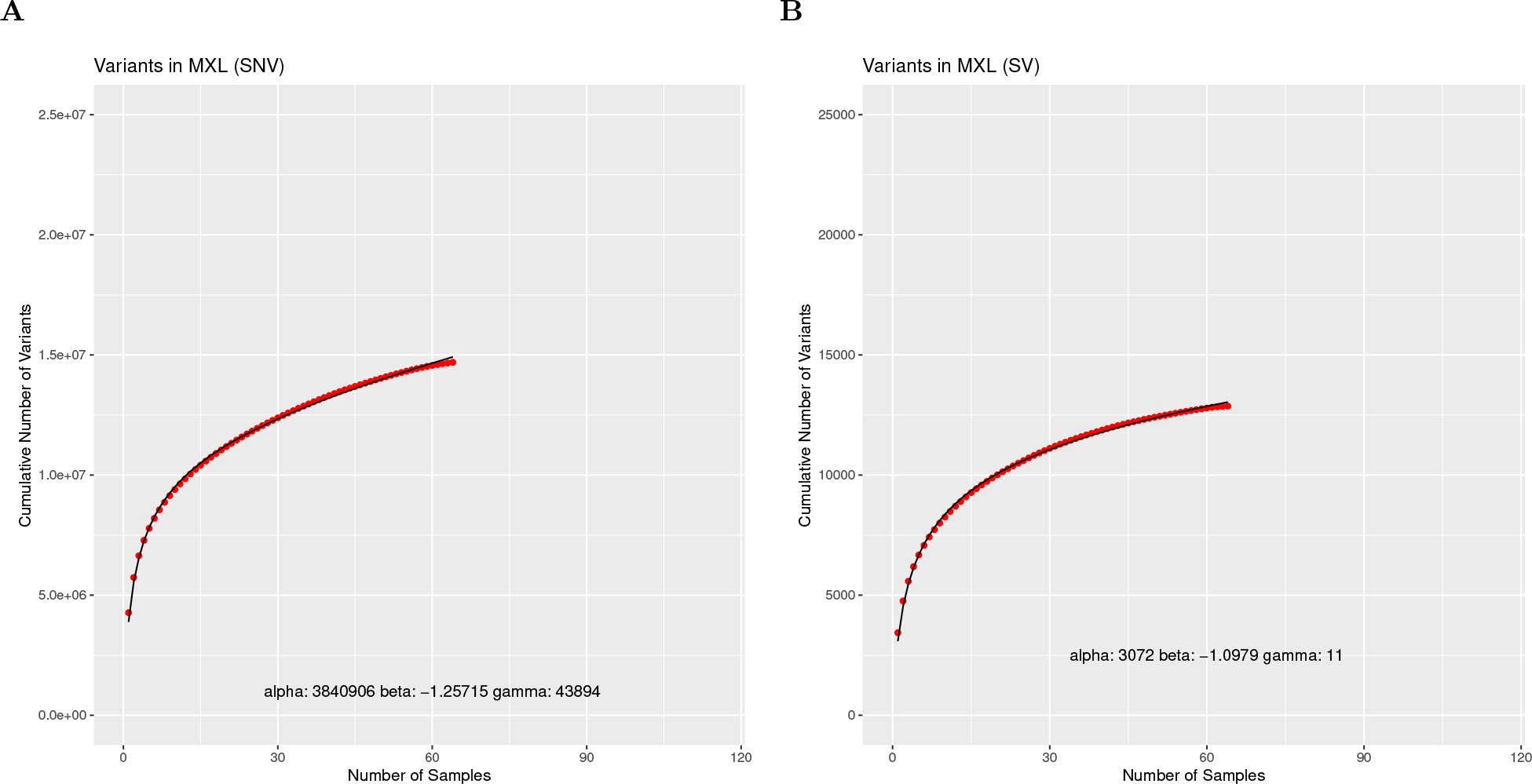
SVCollector curves for the Mexican Ancestry from Los Angeles USA (MXL) population using **(A)** SNV data and **(B)** SV data.

**Supplemental Figure 26:**
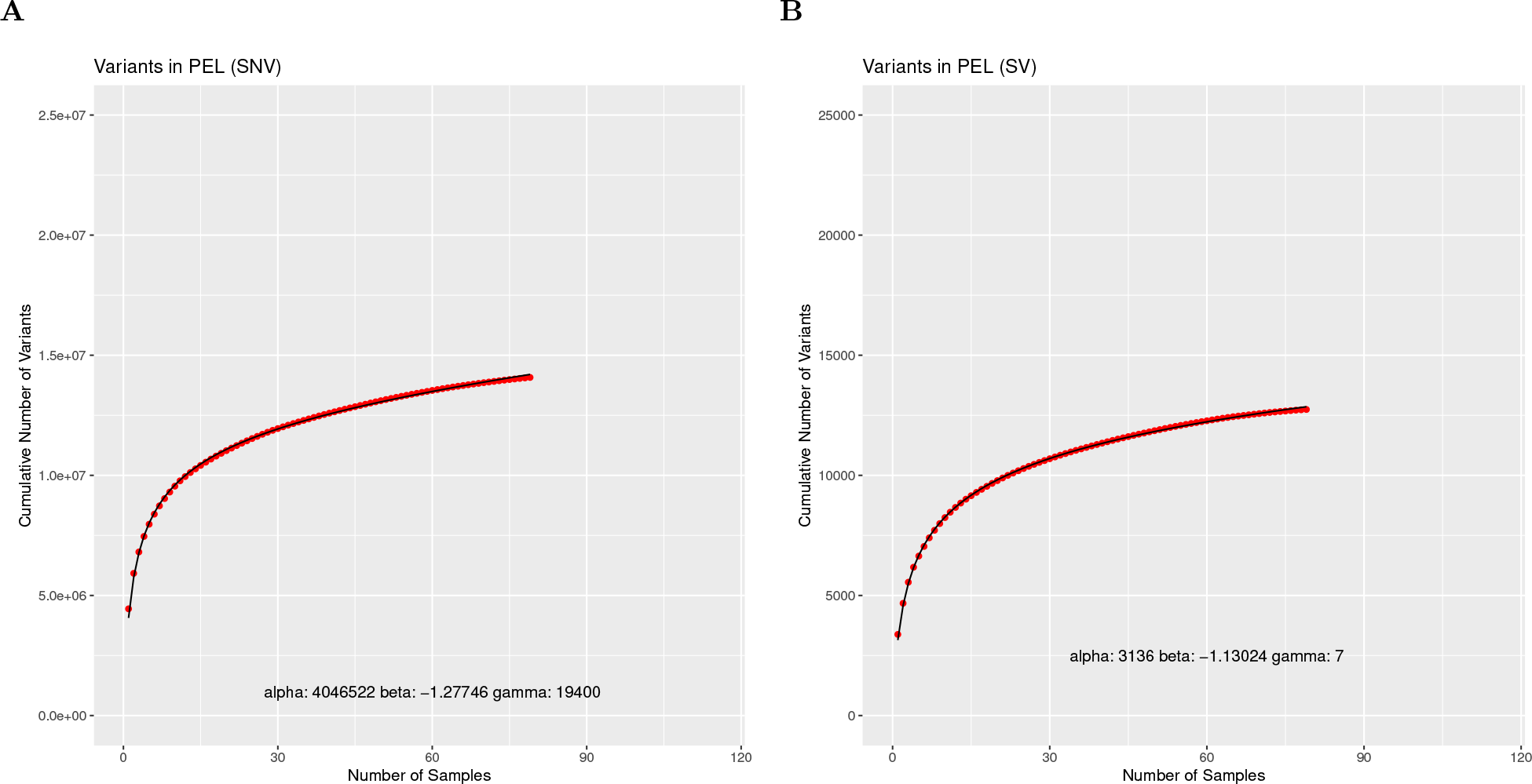
SVCollector curves for the Peruvians from Lima, Peru (PEL) population using **(A)** SNV data and **(B)** SV data.

**Supplemental Figure 27:**
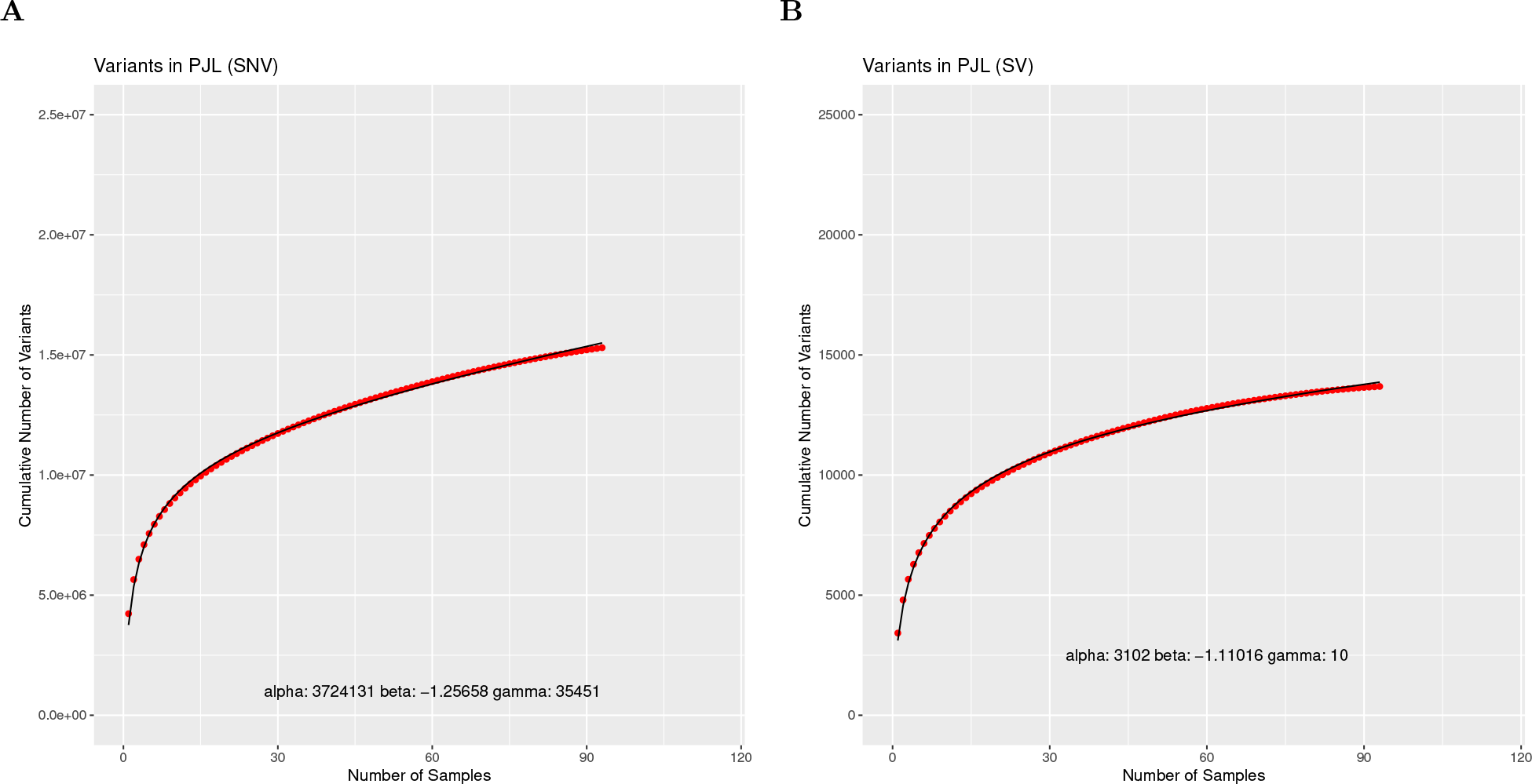
SVCollector curves for the Punjabi from Lahore, Pakistan (PJL) population using **(A)** SNV data and **(B)** SV data.

**Supplemental Figure 28:**
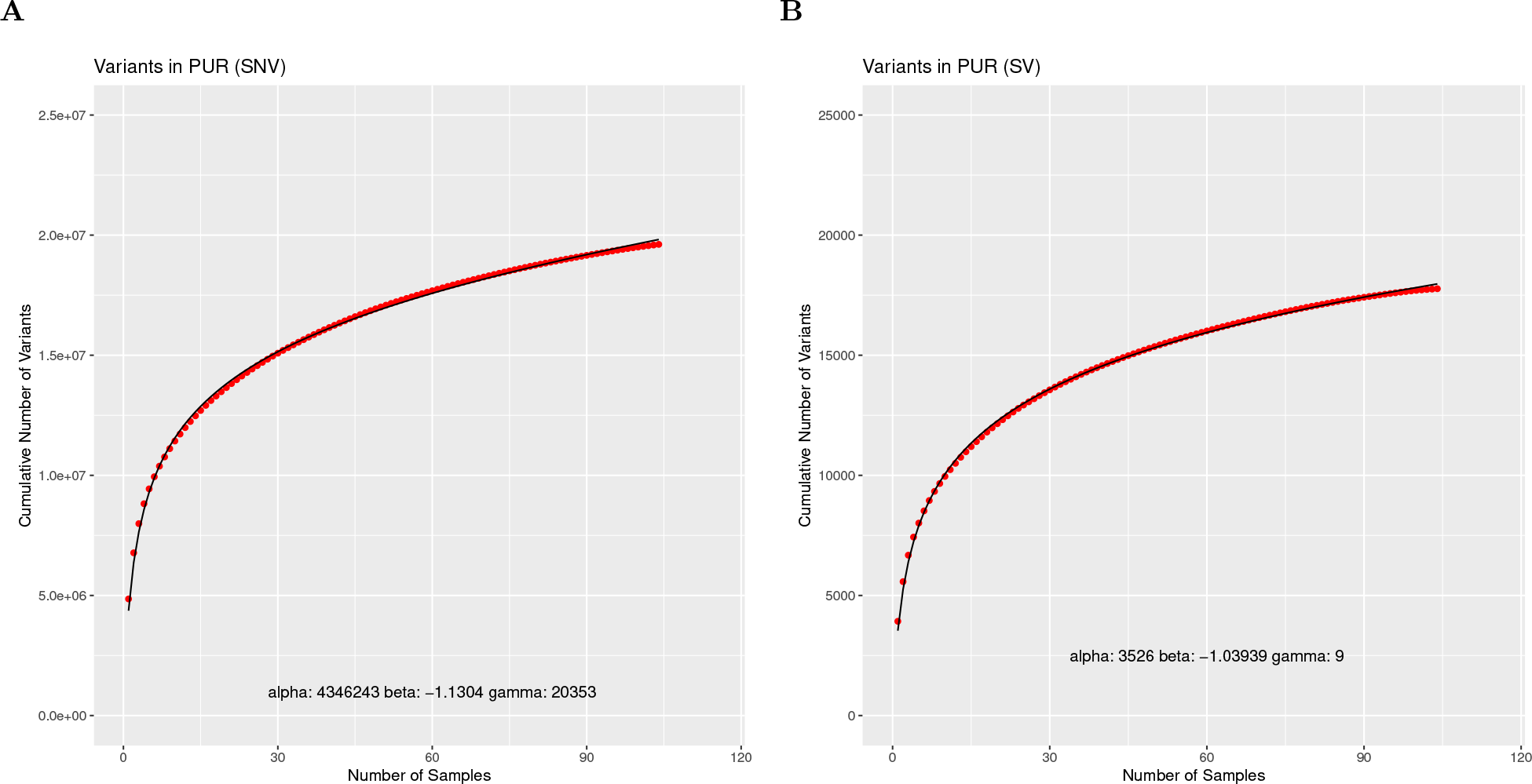
SVCollector curves for the Puerto Ricans from Puerto Rico (PUR) population using **(A)** SNV data and **(B)** SV data.

**Supplemental Figure 29:**
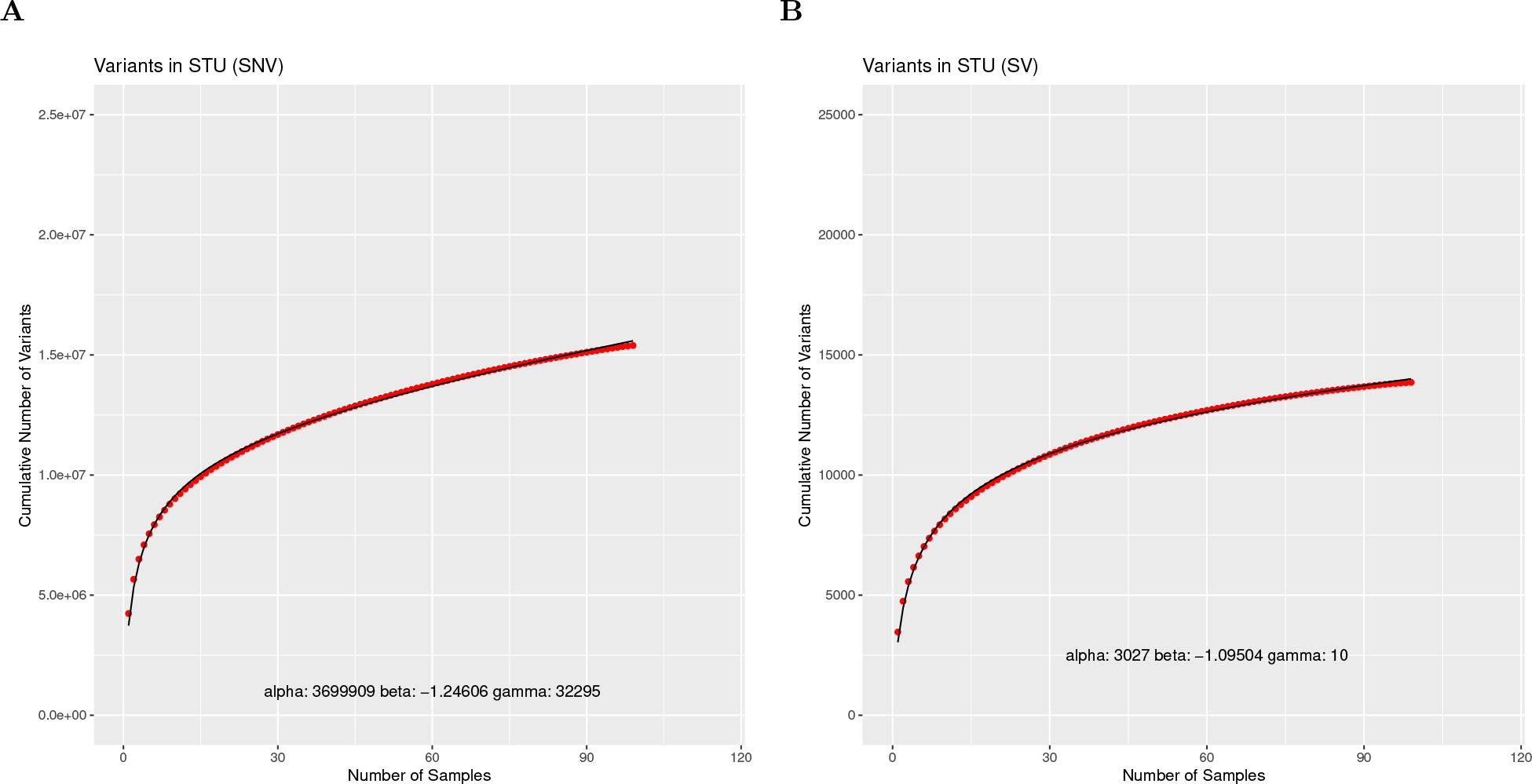
SVCollector curves for the Sri Lankan Tamil from the UK (STU) population using **(A)** SNV data and **(B)** SV data.

**Supplemental Figure 30:**
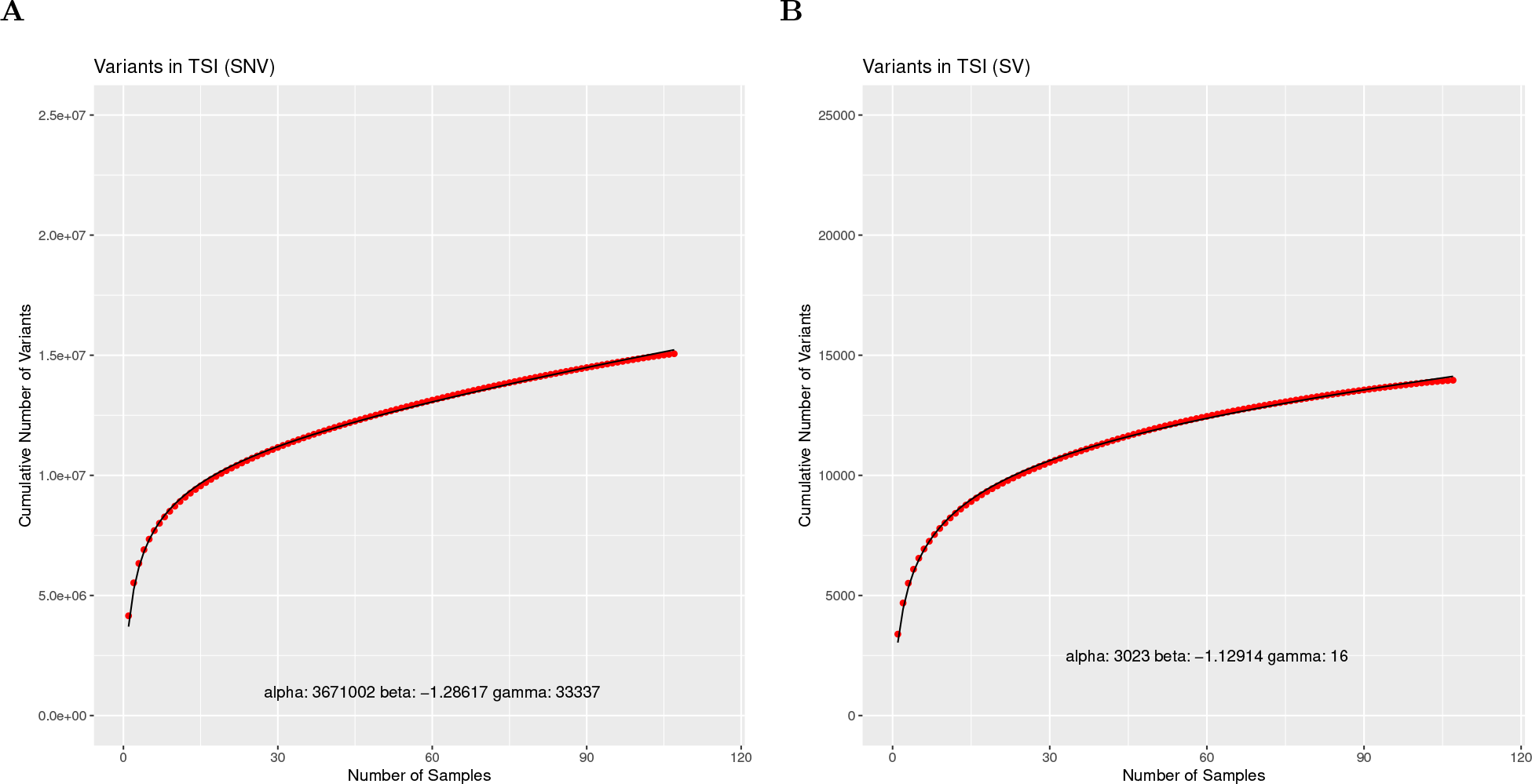
SVCollector curves for the Toscani in Italia (TSI) population using **(A)** SNV data and **(B)** SV data.

**Supplemental Figure 31:**
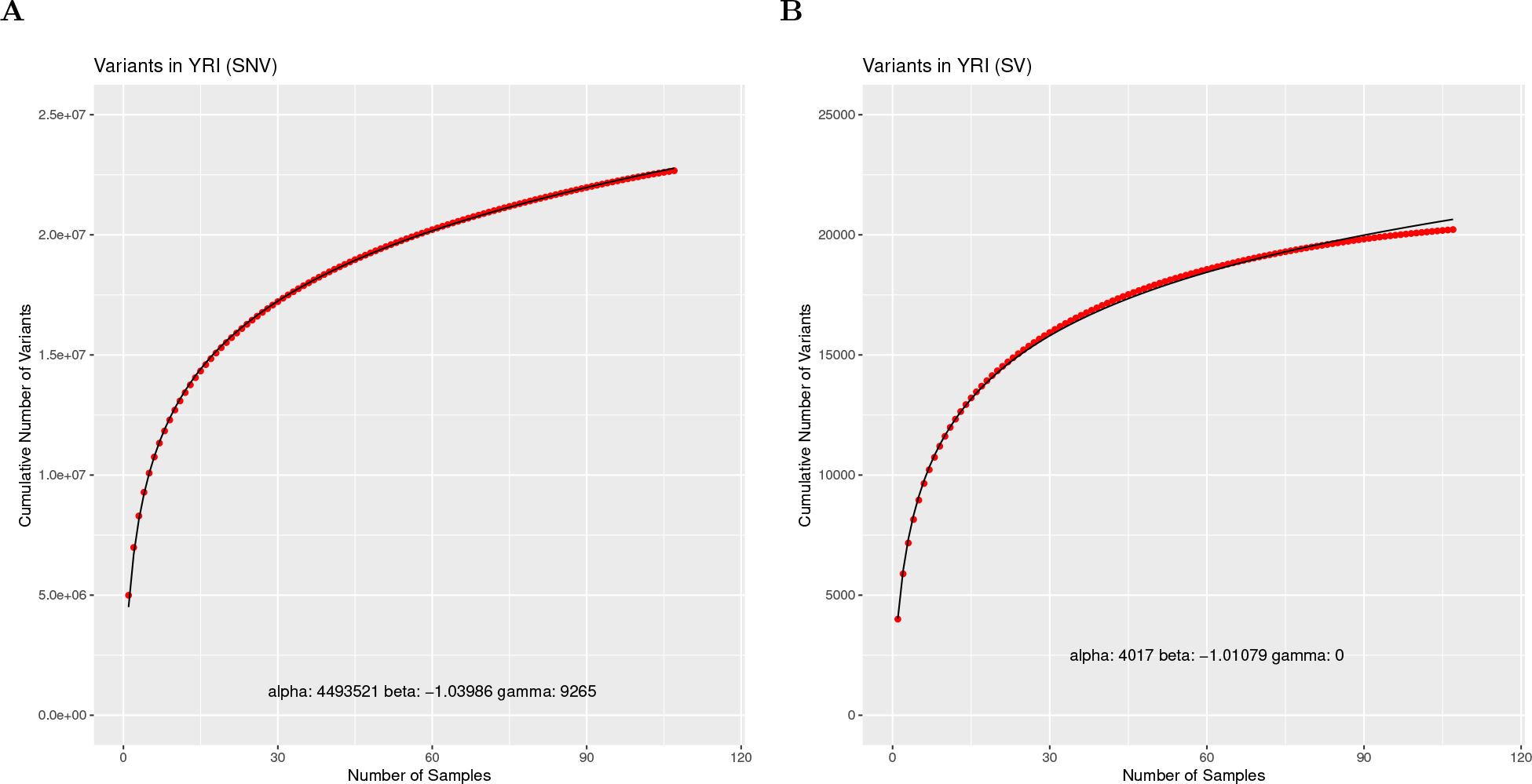
SVCollector curves for the Yoruba in Ibadan, Nigeria (YRI) population using **(A)** SNV data and **(B)** SV data.

